# Dietary Availability Acutely Governs Puberty Onset via Hypothalamic Neural Circuit

**DOI:** 10.1101/2023.09.15.558025

**Authors:** Teppei Goto, Mitsue Hagihara, Satsuki Irie, Takaya Abe, Hiroshi Kiyonari, Kazunari Miyamichi

**Affiliations:** Laboratory for Comparative Connectomics, RIKEN Center for Biosystems Dynamics Research, Kobe, Hyogo 650-0047, Japan; Laboratory for Animal Resources and Genetic Engineering, RIKEN Center for Biosystems Dynamics Research, Kobe, Hyogo 650-0047, Japan

## Abstract

Reproduction poses a substantial burden, especially for mammalian females. Puberty onset serves as a vital checkpoint, regulated based on the body’s energy state, to prevent inappropriate reproductive activity under malnutrition. However, the neural basis of this puberty checkpoint remains poorly understood. Here, we demonstrate that peripubertal malnutrition in female mice reduces the synchronous activity episodes of arcuate kisspeptin neurons (SEs^kiss^), which are critical regulators of the gonadotropin axis. Improved dietary availability boosts SEs^kiss^ frequency, facilitating puberty onset. Using a viral-genetic approach, we show that the activating agouti-related protein neurons in the arcuate nucleus (ARC^Agrp^ neurons), a hunger center, suppresses SEs^kiss^, even with enough food. Conversely, loss-of-function of ARC^Agrp^ neurons enhances SEs^kiss^ during malnutrition, partly promoting irregular sexual maturation. Collectively, a neural circuit connecting feeding to reproductive centers is responsible for disinhibiting SEs^kiss^ frequency based on dietary availability, which sheds light on the neural basis of puberty checkpoint.

**Highlights:**

- The pulsatile activity of arcuate kisspeptin neurons emerges and grows before vaginal opening.
- Pubertal impairment resulting from malnutrition is associated with a reduction in the pulsatile activity of kisspeptin neurons.
- Pubertal recovery by food availability follows the elevated pulsatile activity of kisspeptin neurons during catch-up growth.
- The arcuate Agrp neurons suppress the frequency of pulsatile activity of kisspeptin neurons under negative energy balance.

## Introduction

An adequate, balanced diet is essential for optimal growth and the timely onset of puberty, a critical stage for acquiring reproductive capacity. The puberty onset is intricately tied to the body’s energy balance. Contemporary women experience earlier puberty compared with previous generations, possibly due to excessive consumption of high-fat, processed foods^1^. Conversely, anorexia nervosa rates have risen among females under the age of 15 years^2^, linked to delayed puberty and compromised reproductive health in adulthood. In female rodent models, overnutrition leads to increased body weight and early puberty, while chronic energy deficiency, such as that induced by malnutrition, delays puberty^3,4^. However, our understanding of the neural circuits that mediate nutritional and metabolic signals’ effects on reproductive centers remains limited.

The regulation of the reproductive function of adult females primarily hinges on the gonadotropin-releasing hormone (GnRH) produced by neurons in the anterior hypothalamus^5^. GnRH is released in a pulsatile manner, with intervals ranging from 30 minutes to several hours, varying by species and reproductive cycle^6^. This pulsatile secretion is crucial for reproductive functions^7^. Pulsatile GnRH secretion is mainly governed by kisspeptin neurons in the arcuate nucleus (ARC^kiss^ neurons), acting as the GnRH pulse generator^8–10^. Indeed, mice and rats with deficient *Kiss1* gene, which encodes kisspeptin, are infertile^11–13^. This has led to the prevailing view that the onset of puberty is driven by the emergence of GnRH pulse generator activity in the ARC^kiss^ neurons.

If ARC^kiss^ neurons play a pivotal role in the regulation of puberty onset, how do they perceive and respond to nutritional and metabolic cues originating from the body? One plausible explanation involves the humoral factors^4,14^, including metabolites and hormones, likely through mechanisms that operate outside the blood-brain barrier. For instance, leptin, derived from adipose tissue, is implicated, as evidenced by the prevention of puberty onset in both whole-body and brain-specific *leptin receptor* knockout models^15,16^. However, it is important to note that genetic mutant models have mainly explored extreme leptin signaling scenarios, which may be less attainable under physiological dietary conditions. Consequently, the extent to which hyper- or hypo-leptinemia influences nutrition-associated puberty timing may vary with experimental paradigms^17–19^. That being said, previous research has suggested potential neural pathways for transmitting leptin signaling into the reproductive circuitry. Agouti-related protein-expressing neurons in the arcuate nucleus (ARC^Agrp^ neurons), which serves as a well-established hunger center^20^, are implicated in the leptin-mediated control of puberty timing^21^, alongside other contributing neurons^22^. These ARC^Agrp^ neurons are known to form mono-synaptic connections to ARC^kiss^ neurons and may exert a negative influence on adult female reproductive functions^23^, making the ARC^Agrp^-to-ARC^kiss^ circuitry an intriguing subject warranting further investigation, although its role in puberty onset remains poorly understood.

Another line of evidence suggests the involvement of cellular energy sensors capable of epigenetically regulating *Kiss1* mRNA expression within ARC^kiss^ neurons in response to the body’s metabolic status^24,25^. Previous studies have emphasized the importance of the mammalian target of rapamycin (mTOR)^18^, Sirtuins^26^, and the AMP-activated protein kinase^27^ within ARC^kiss^ neurons. However, due to the robust regulation of energy homeostasis, these molecular sensing systems exhibit a slow response, potentially integrating information over the course of days to weeks. Given that puberty timing is particularly susceptible to alterations in nutritional cues and energy reserves^3,4^, we can postulate the existence of more rapidly acting mechanisms, although these have not yet been elucidated.

To comprehensively understand the regulatory mechanisms governing puberty onset, direct visualization of neural activity dynamics of ARC^kiss^ neurons is crucial. Previously, assessing GnRH pulse generators relied on repetitive measurements of circulating levels of gonadotorophin^28^, a procedure that can induce stress in young animals. Recent research has achieved cell-type-specific, chronic Ca^2+^ imaging of ARC^kiss^ neuron activities in freely moving mice using fiber photometry^8,9,29,30^. In adult mice, ARC^kiss^ neurons display pulsatile activities, termed synchronous episodes of elevated Ca^2+^ (referred to as SEs^kiss^ for simplicity). SEs^kiss^ are closely tied to gonadotropin secretion^9^ and regulated by sex steroid hormones^31^. SEs^kiss^ frequency varies with the estrus cycle^8,30^. However, the system’s applicability to studying puberty onset remains uncertain. Hence, our goal is to employ fiber photometry to characterize the activity dynamics of ARC^kiss^ neurons under various dietary conditions and unravel the neural circuitry responsible for transmitting feeding-related signals to the ARC^kiss^ neurons during the peripubertal stage in female mice.

## Results

### SEs^kiss^ emerge before the vaginal opening

We first aimed to chronically visualize the activities of ARC^kiss^ neurons during the peripubertal stage of normally nourished female mice. To this aim, we crossed *Kiss1-Cre* mice^32^ with a Cre-dependent mouse line driving GCaMP6s Ca^2+^ sensor^33,34^ to obtain double heterozygous female mice. Using these mice, we employed fiber photometry imaging of ARC from postnatal day (PND) 24 to 45 (Figure 1A). The location and specificity of GcaMP6s-positive cells were verified through histochemistry (Figure 1B–D). Body weight was unaltered irrespective of genotype or surgical intervention (Figure 1E). Two Ca^2+^ imaging sessions were conducted every third day, each lasting 6 hours, in both light and dark periods. Representative photometric signals showed sharp pulsatile activities of ARC^kiss^ neurons that highly resembled SEs^kiss^ detected in the adult mice^8,9,30^ around PND 24–30 (Figure 1F). Raster plot representation revealed that SEs^kiss^ were detected in all four female mice we examined before the occurrence of vaginal opening, the first sign towards puberty onset (Figure 1G, H). The number and intensity of SEs^kiss^ increased with age, plateauing at PND 42 and 35, respectively (Figure 1I). SEs^kiss^ were consistent between light and dark periods (Figure 1J), in contrast to the nocturnal elevation observed in the gonadotropin axis of primates^35–37^. Collectively, these results reveal that SEs^kiss^ appear substantially before the onset of puberty, consistent with their roles in promoting puberty initiation^9,38,39^.

**Figure 1.**
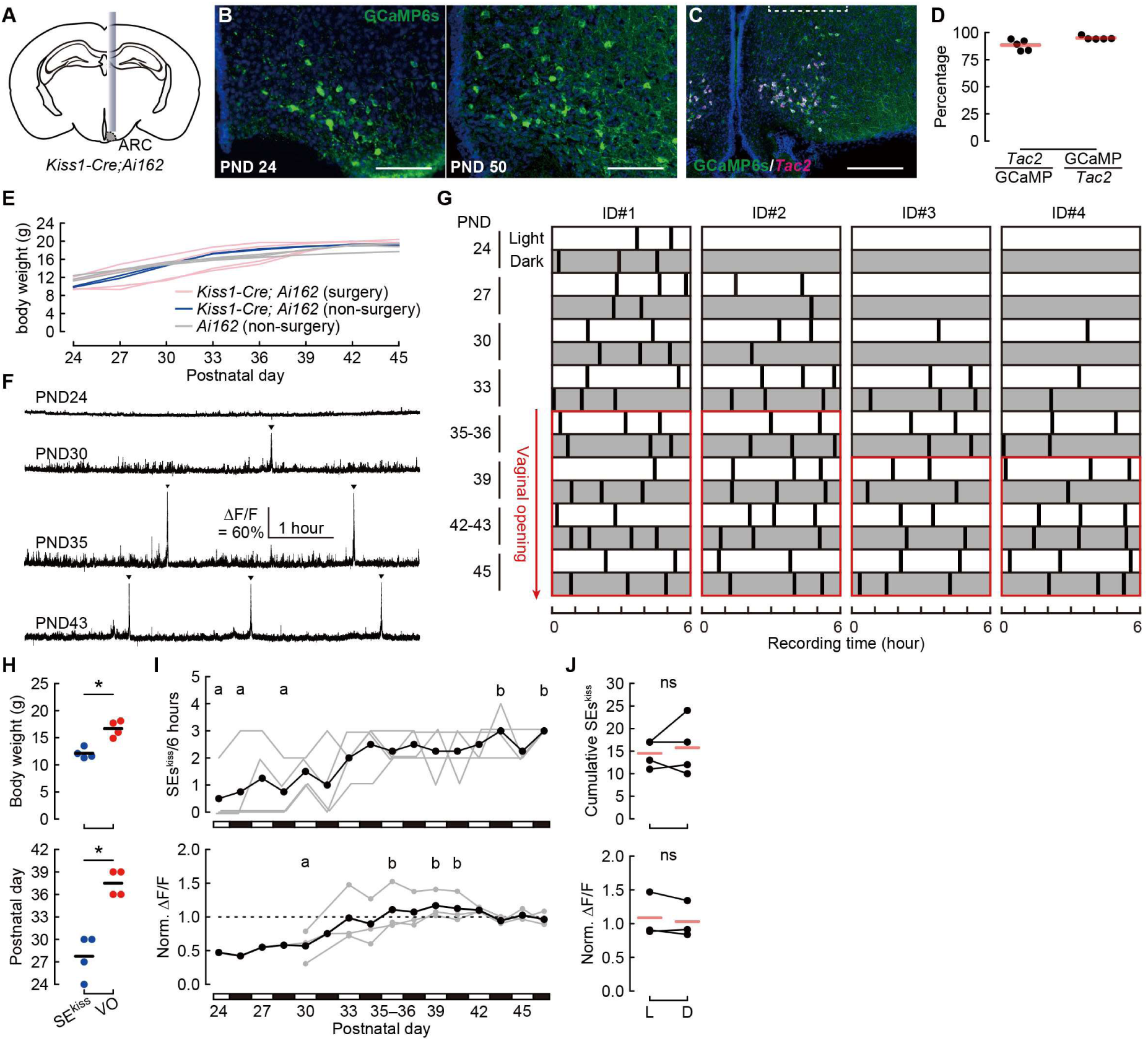
Appearance of SEs^kiss^ before vaginal opening. (A) Schematic of the experimental setting. (B) Representative coronal sections of the ARC from *Kiss1-Cre; Ai162* double heterozygous mice showing GCaMP6s expression (stained by anti-green fluorescent protein [GFP]) at postnatal day (PND) 24 and 50. Scale bars, 100 μm. (C) Representative coronal section of the ARC from the *Kiss1-Cre; Ai162* female mice showing optic fiber tract (dotted line) with *Tac2* mRNA expression (magenta, a marker of ARC^kiss^ neurons) and GCaMP6s (green, stained by anti-GFP) counterstained with DAPI (blue). Scale bar, 200 μm. (D) Quantification of specificity (*Tac2*+/GCaMP6s+) and efficiency (GCaMP6s+/*Tac2*+), n = 5. (E) Growth curves of *Kiss1-Cre; Ai162* mice with or without optic fiber implantation (surgery) and control mice with only the *Ai162* allele without surgery. (F) Representative 6-hour photometry traces showing SEs^kiss^ (arrowheads) during the light period at PND 24, 30, 35, and 43. (G) Raster plots of SEs^kiss^ (vertical bars) during the peripubertal stages. Open and gray boxes indicate the light and dark periods, respectively. Red frames indicate the timings after vaginal opening (VO). (H) Body weight (upper) and PND (lower) at the first observation of SEs^kiss^ and VO. *, p < 0.05 by two-sided paired *t*-test. Notably, body weight at the first emergence of SEs^kiss^ was significantly lower than that at the timing of VO. (I) Average number (upper) and intensity (lower) of SEs^kiss^ per 6-hour time window during peripubertal stages. Individual data are shown in gray. Different letters (a and b) denote significant differences at p < 0.05 by one-way ANOVA with repeated measures followed by the Tukey–Kramer post-hoc test. Note that ID #3 was excluded from the intensity analysis because the patch code was changed during the experiment. (J) Top: Total number of SEs^kiss^ during the light (L) or dark (D) period observed in the 48-hour imaging sessions (6 hours per day × 8 days, as shown in panel G). Bottom: Normalized change in fluorescent intensity (ΔF/F) during the light or dark periods among PND 33–39. ns, no significant difference by two-sided paired *t*-test. Intensity was normalized within individuals to the average intensity between PND 42–43 and 45 (I, J). n=4 (G–J).

### Dynamic regulation of SEs^kiss^ by energy balance

We next aimed to visualize the peripubertal activity dynamics of ARC^kiss^ neurons under various feeding conditions. To this end, we validated the protocol for the food restriction-induced impairment of sexual maturation^25,27,40^ as defined by the vaginal opening and first estrus via vaginal cytology^41,42^. We also examined the potential effects of exogenous supplementation of leptin (Figure 2A). Wild-type female mice were housed under four distinct conditions: i) *ad libitum* food access (AL); ii) food restriction (FR) to 2.5 grams per day (representing approximately a 30% reduction in daily caloric intake; Figure S1A–F) provided in a single administration during a light period; iii) FR with chronic leptin supplementation (FR+leptin) administered via an osmotic minipump, and iv) FR→AL in the interim at PND 35.

**Figure 2.**
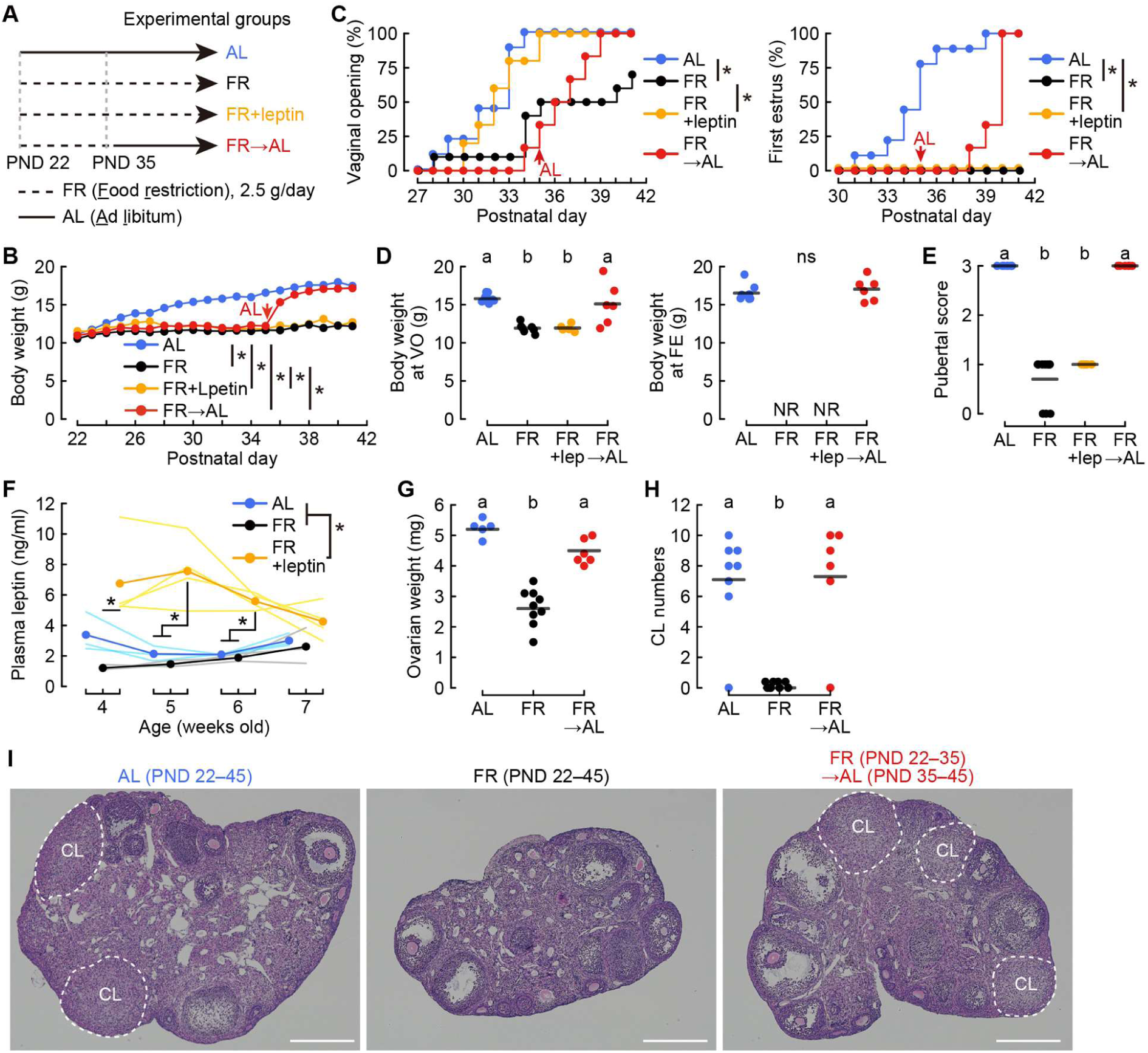
Puberty onset under various food conditions and leptin supplementation. (A) Timetable of feeding conditions and leptin administration. PND, postnatal day. (B, C) Growth curve (B) and cumulative probability of puberty onset, denoted by vaginal opening (C, left) and vaginal cytological first estrus (C, right) of the peripubertal mice under the AL (blue), FR (black), FR with exogenous leptin administration (yellow), and FR→AL (red, switching at PND 35 as shown by the red arrow) conditions. *, p < 0.05 by the Kolmogorov–Smirnov test with Bonferroni correction. (D, E) Body weight at the time of vaginal opening (D, left) and first estrus (D, right), and pubertal score (E) in the AL, FR, FR+leptin, and FR→AL groups. The mice were assessed on a 0–3 scale according to their vaginal state: absence of vaginal opening (score 0); the presence of vaginal opening and diestrus (1), proestrus (2), and estrus (3). Different letters (a, b) denote a significant difference (p < 0.05) by one-way ANOVA followed by the Tukey–Kramer post-hoc test (D, left) or Kruskal–Wallis test followed by the Mann– Whitney *U* test (E). ns, not significant (D, right) by the Mann–Whitney *U* test. NR, no record. n = 9 for AL, n = 10 for FR, n = 5 for FR+leptin, and n = 6 for the FR→AL groups (B–E). (F) Averaged plasma leptin levels in the AL (light blue), FR (black), and FR+leptin (yellow) groups, with light color lines representing individual raw data. Two-way ANOVA with repeated measures; condition effect, p < 0.01; time course effect, ns; interaction effect, p < 0.05. *, p < 0.05 by one-way ANOVA followed by the Tukey–Kramer post-hoc test. n = 3 for AL and FR and n = 4 for FR+leptin conditions. (G, H) Ovarian weight (G) and number of corpus lutea (CL) (H) at PND 45. Letters (a and b) denote a significant difference (p < 0.05) by one-way ANOVA followed by the Tukey–Kramer post-hoc test (G), or the Kruskal–Wallis test followed by the Mann– Whitney *U* test (H). n = 5 or 8 for AL, 9 for FR, and n = 6 for FR→AL conditions. (I) Representative images of hematoxylin/eosin-stained ovarian sections from the AL, FR, and FR→AL groups at PND 45. CL, corpus luteum. Scale bars, 200 μm. For more data, see Figure S1.

Under the AL condition, all female mice exhibited vaginal opening by PND 35 and first estrus by PND 39. Under the FR condition, the female mice did not surpass a body weight of 13 grams by PND 42, only 50% demonstrated vaginal opening, and none exhibited first vaginal estrus (Figure 2B–D). Notably, the growth curve was not changed by leptin supplementation to the FR group (Figure 2B). Chronic leptin supplementation facilitated vaginal opening but failed to alleviate the absence of estrus under the FR condition (Figure 2C). Body weight at the vaginal opening was significantly lower in the FR and FR+leptin groups compared with the AL group (Figure 2D). The pubertal score defined by the progress of puberty onset (Method) was also significantly lower in the FR and FR+leptin groups (Figure 2E). We showed that plasma leptin level was continuously and significantly higher in the FR+leptin group, whereas only modestly lower in the FR group, compared with the AL control (Figure 2F). Collectively, these data suggest that, under our FR condition, hypoleptinemia cannot explain the FR-induced impairment of sexual maturation based on the first vaginal estrus, despite the potential of leptin as a puberty-promoting hormone^15,21^.

Upon removal of the FR at PND 35 (FR→AL group), all mice showed catch-up growth (Figure 2B) and reached the vaginal first estrus within 3–5 days (Figure 2C, D). Mice displaying sexual maturation (AL and FR→AL groups) exhibited a significantly higher pubertal score (Figure 2E). To examine the ovarian phenotype in these mice, we harvested ovaries at PND 45 and analyzed images of hematoxylin/eosin (H/E)-stained ovarian sections. At PND 45, Mice displaying sexual maturation (AL and FR→AL groups) had a heavier ovary with the formation of a postovulatory corpus luteum (CL), whereas no CL formation was observed in the FR group (Figure 2G–I). These data confirmed that the puberty impairment caused by FR can be reversed through refeeding^43,44^ in a manner partially independent of leptin signaling^17^ (however, also see ref.^18^).

Based on these findings, we decided to focus on SEs^kiss^ at PND 37–40 under the chronic FR and FR→AL conditions by fiber photometry. To target the ARC^kiss^ neurons selectively and efficiently, we employed a Cre-dependent adeno-associated virus (AAV) that drives GCaMP6s into the ARC of *Kiss1*-*Cre* female mice (Figure 3A), rather than a low-throughput double-transgenic strategy (Figure 1). Post-hoc histochemical analyses confirmed the fiber location and the efficiency and specificity of GCaMP6s expression (Figure 3B, C). Two Ca^2+^ imaging sessions were conducted daily, each lasting 6 hours, in both light and dark periods. During the FR conditions, we provided 2.5 grams of food soon after the start of the Ca^2+^ imaging session in the light period. At PND 37 of the FR condition, although the female mice did not surpass a body weight of 13 grams (Figure 3D), we detected noticeable SEs^kiss^ (Figure 3E, F), albeit at a low frequency of approximately 0.3 per hour. In addition, the waveform of SEs^kiss^ in the dark periods (> 12 hours after food supply) was less stereotypical (Figure S2A, B). These data suggest that SEs^kiss^ can be formed even when the body’s energy state impedes sexual maturation, and that food availability can influence the frequency and waveform of SEs^kiss^.

**Figure 3.**
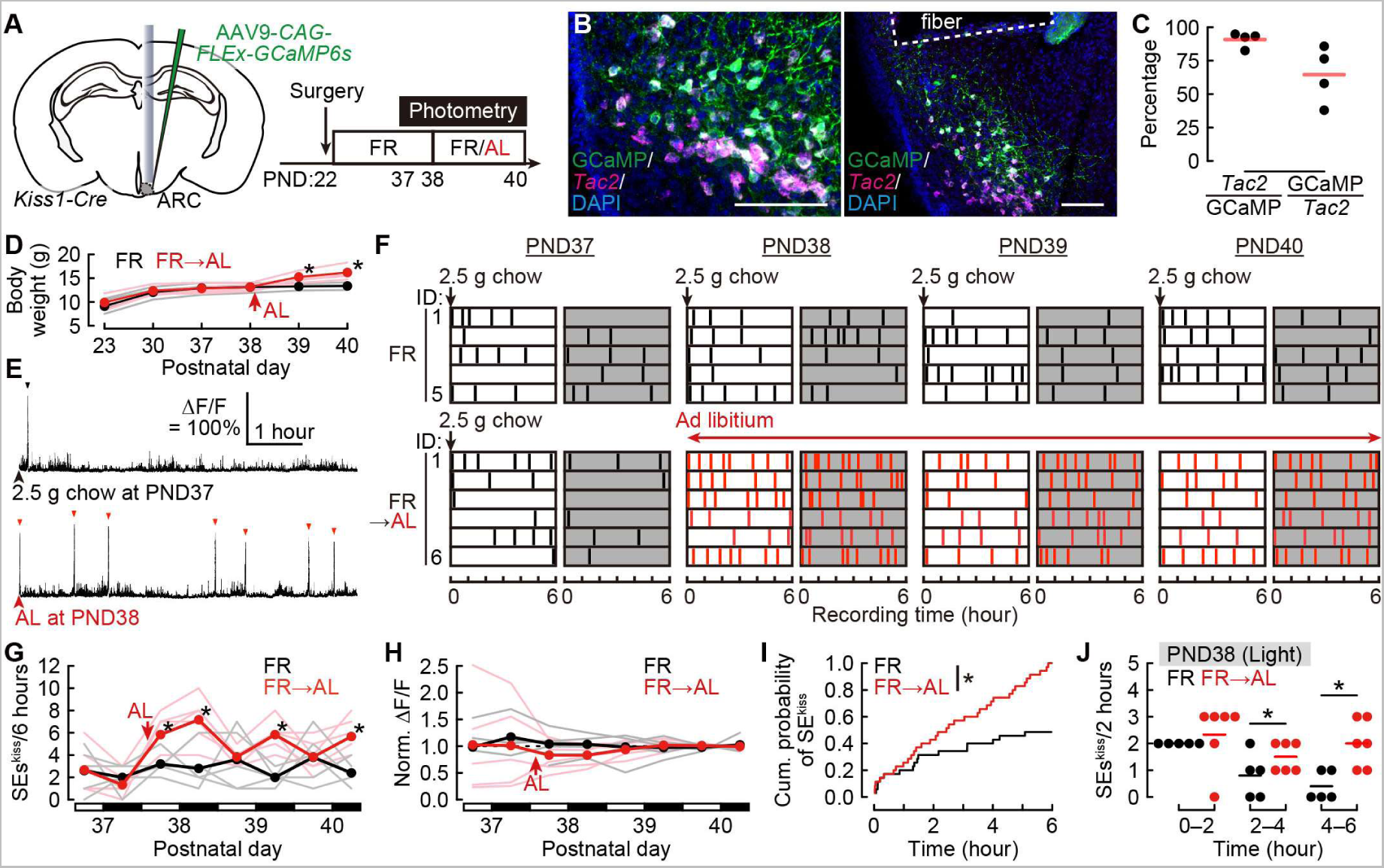
Dynamic modulation of peripubertal SEs^kiss^ by changing energy balance. (A) Schematics of the experimental setup and time line. (B) Representative coronal brain sections of the ARC from *Kiss1-Cre* female mice showing optic fiber tract with *Tac2* mRNA expression (magenta, a marker of ARC^kiss^ neurons) and GCaMP6s (green, stained by anti-GFP) counterstained with DAPI (blue). Scale bars, 100 μm. (C) Quantification of specificity (*Tac2*+/GCaMP6s+) and efficiency (GCaMP6s+/*Tac2*+). n = 4. (D) Body weight under the FR and FR→AL (switching at PND 38 as shown by the red arrow) conditions. Two-way ANOVA with repeated measures; feeding condition effect, ns; PND effect, p < 0.01; interaction effect, p < 0.01. *, p < 0.01 by post-hoc two-tailed unpaired *t*-test. (E) Representative photometry traces showing SEs^kiss^ (arrowheads) in the light period under the FR (at PND 37) and soon after switching to AL at PND 38. (F) Raster plots of SEs^kiss^ (vertical bars) under the FR and FR→AL conditions. Open and gray boxes show light and dark periods, respectively. (G, H) Number (G) and infrared intensity by normalized (norm) ΔF/F (H) of SEs^kiss^ at 6-hour windows under both conditions. Two-way ANOVA with repeated measures; feeding condition effect, p < 0.05; period effect, p < 0.01; interaction effect, p < 0.01 in SEs^kiss^ number (G). *, p < 0.05 by post-hoc two-tailed unpaired *t*-test. ns in SEs^kiss^ intensity (H) by two-way ANOVA with repeated measures. (I) Cumulative probability of SEs^kiss^ under the FR (black) and FR→AL (red) conditions in the light period at PND 38. *, p < 0.05 by the Kolmogorov–Smirnov test. (J) Number of SEs^kiss^ within a 2-hour time window under FR (black) and soon after the removal of FR (red). Two-way ANOVA with repeated measures; feeding condition effect, p < 0.05; time course effect, p < 0.01; interaction effect, ns. *, p < 0.05 by two-tailed unpaired *t*-test with Bonferroni correction. n = 5 for the FR and n = 6 for the FR→AL condition (D–J). For more data, see Figure S2.

Upon removal of the FR at PND38, the frequency of SEs^kiss^ was significantly increased without any alterations in amplitude (Figure 3G, H). The variability of the SEs^kiss^ waveform was also restored (Figure S2A, B). A notably higher frequency was observed during the dark period (Figure 3F, G), indicating a potential correlation with nocturnal excessive feeding during catch-up growth. A higher temporal resolution analysis revealed that the chronic FR and FR→AL conditions diverged 4–6 hours after the removal of the FR (Figure 3I, J); mice subjected to FR displayed a declining trend of SEs^kiss^ over time following the supply of 2.5 grams of food, whereas mice subjected to FR→AL sustained elevated frequency of SEs^kiss^. Notably, the dynamics of blood glucose levels peaked 10 minutes after food supply and gradually declined thereafter (Figure S2C), which contrasts with the temporal frequency patterns of SEs^kiss^. Together, these observations suggest that the recuperation from FR is characterized by a prolonged increased frequency of SEs^kiss^. This phenomenon occurs within several hours after the amelioration of dietary availability, well in advance of the vaginal estrus event.

### Dual-color recording of ARC^Agrp^ and ARC^kiss^ neurons

To understand how the activity of ARC^kiss^ neurons is modulated by dietary availability, we investigated the possibility of a monosynaptic inhibitory input to the ARC^kiss^ neurons that is active during starvation and suppressed by refeeding. The ARC^Agrp^ neurons possess the desired characteristics of being GABAergic and monosynaptically connecting to ARC^kiss^ neurons^23,45^. ARC^Agrp^ neurons are known to be active during fasting and are rapidly suppressed upon exposure to food-related stimuli^46–48^. However, activity dynamics of ARC^Agrp^ neurons in malnourished female mice during the peripubertal stage remain unknown. To gain genetic access to ARC^Agrp^ neurons independent of ARC^kiss^ neurons, we utilized CRISPR-mediated genome editing^49^ to generate *Agrp-Flpo* knock-in mice (Figure S3).

To investigate the relationship between the activities of ARC^Agrp^ and ARC^kiss^ neurons, we employed dual-color Ca^2+^ imaging in PND 37–38 *Kiss1-Cre; Agrp-Flpo* double heterozygous female mice under the FR→AL condition (Figure 4A). We selectively targeted GCaMP6s to ARC^Agrp^ neurons and jRCaMP1a^50^ to ARC^kiss^ neurons. Post-hoc histochemical analyses revealed the high specificity of both targeting methods, with no leaky expressions observed in nontargeted cell types (Figure 4B, C). We were also able to monitor GCaMP6s and jRCaMP1a signals simultaneously without interference (Figure S4): When both sensors were used to detect SEs^kiss^, we confirmed that all peaks detected with GCaMP6s were also detected with jRCaMP1a.

**Figure 4.**
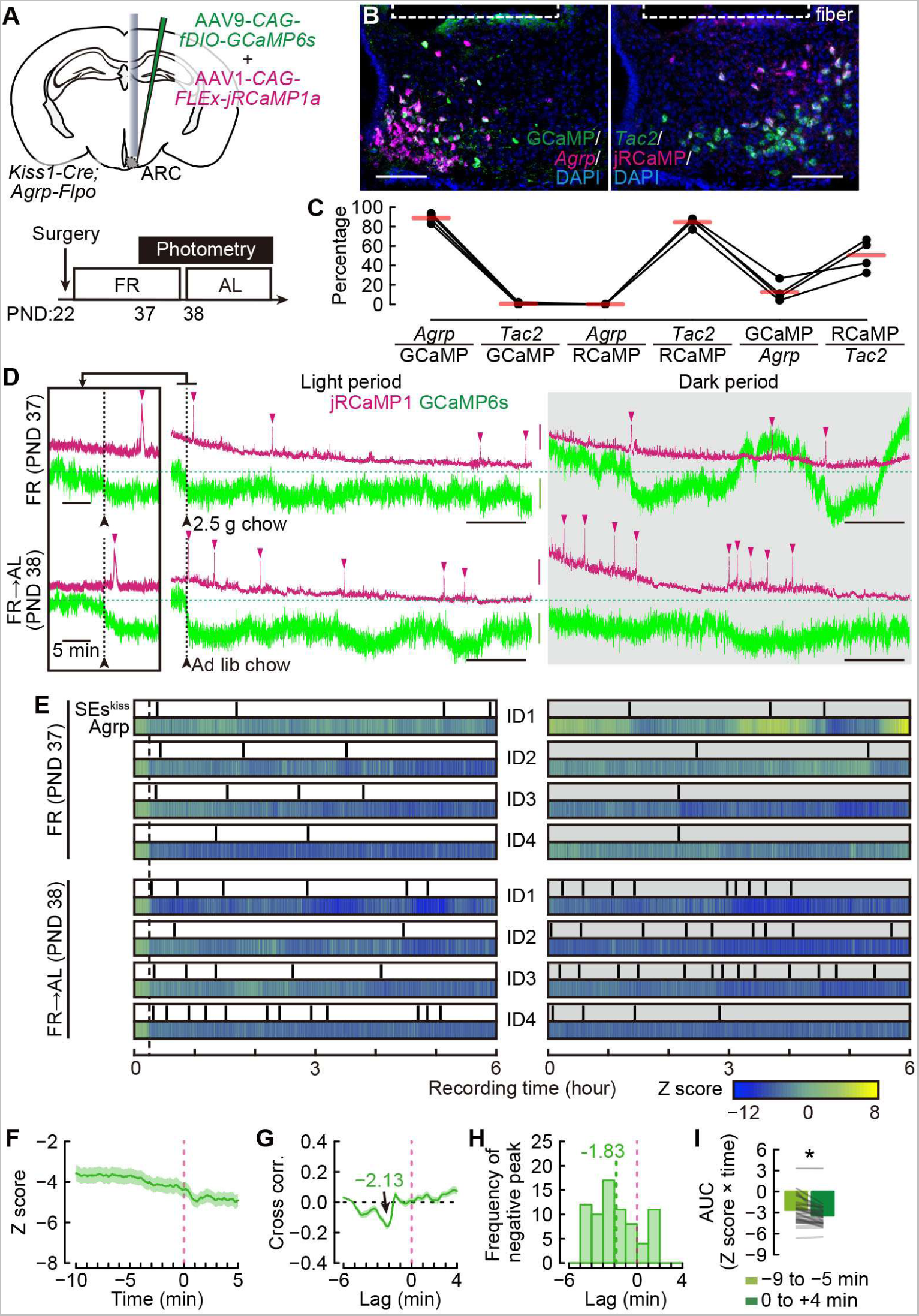
Peripubertal dual-color recording of ARC^Agrp^ and ARC^kiss^ neurons. (A) Schematic of the experimental setup and time line. (B) Representative coronal sections of the ARC showing *Agrp* mRNA expression (magenta), GCaMP6s (green) (left) and *Tac2* mRNA expression (green, a marker of ARC^kiss^ neurons), and jRCaMP1a (magenta) (right) counterstained with DAPI (blue). Dotted lines represent the optic fiber tract. Scale bars, 100 μm. (C) Quantification of targeting specificities (*Agrp*+/GCaMP6s+ and *Tac2*+/jRCaMP1a+), ratios of leaky expression to the untargeted cell type (*Tac2*+/GCaMP6s+ and *Agrp*+/jRCaMP1a+), and targeting efficiencies (GCaMP6s+/*Agrp+* and jRCaMP1a+/*Tac2+*). Of note, the targeting efficiency to ARC^Agrp^ neurons is lower, presumably due to the low efficiency of Flpo-mediated recombination, n = 4. (D) Representative 6-hour dual-color photometry traces showing SEs^kiss^ (magenta arrowheads) under the FR condition and after switching to the AL condition. Green, GCaMP6s signals. Magenta, jRCaMP1a signals. Horizontal black scale bars, 1 hour. Vertical green and magenta scale bars, ΔF/F 5%. The left boxes show higher temporal magnification around the food supply (dotted vertical lines and black arrowheads). (E) Raster plots of SEs^kiss^ (vertical bars) and heatmap showing Z-scored photometric signals of ARC^Agrp^ neurons under FR condition (top) and following the removal of FR (bottom). The left and right panels show the results for the light and dark periods, respectively, n = 4. (F) The averaged Z-scored photometric traces of ARC^Agrp^ neurons (green lines) relative to SEs^kiss^ events (pink dotted lines). (G) Cross-correlation (green lines) between Z-scored photometric signals of ARC^Agrp^ neurons and photometric signals of ARC^kiss^ neurons. 0 is the peak of SEs^kiss^ (pink dotted lines). Average peak lag time = −2.13 minutes. The shadows represent the SEM (F and G). (H) Frequency of negative peaks of the cross-correlation. Average (green dotted line) = –1.83 minutes. (I) Area under the curve values within −9 to −5-minute and 0 to +4-minute time windows. *, p < 0.05 by two-sided Wilcoxon signed-rank test with Bonferroni correction. n = 73 events (F–I). For more data, see Figures S3 and S4 and Movie S1.

Following food supply, we observed an immediate reduction in the photometric signals of the ARC^Agrp^ neurons, consistent with previous findings^46–48^ (Figure 4D, E, and Movie S1). The number of SEs^kiss^ remained low during and increased upon the removal of FR, consistent with Figure 3G. The activities of the ARC^Agrp^ neurons were low for 6 hours following food supply and subsequently displayed considerable fluctuations during the dark period. Notably, instances of spontaneous reduction in the activities of ARC^Agrp^ neurons were frequently associated with SEs^kiss^ events (Figure 4D, E). Indeed, the Z-scored photometric signals of the ARC^Agrp^ neurons showed a substantial decline just before the SEs^kiss^ event (Figure 4F). To further probe the temporal sequence of activity change in ARC^Agrp^ neurons and SEs^kiss^, we analyzed the cross-correlation between the peak of SEs^kiss^ and Z-scored photometric signals of ARC^Agrp^ neurons. If activities of ARC^Agrp^ neurons precede SEs^kiss^ events, the peak of cross-correlation would appear within the temporal expanse of negative time lag. We found negative peaks around −5 to −1 minutes relative to the SEs^kiss^ (Figure 4G, H), suggesting that SEs^kiss^ tended to occur after the reduction in ARC^Agrp^ neuron activity. Quantification of the area under the curve (AUC) of Z-scored photometric signals of ARC^Agrp^ neurons also supported this notion (Figure 4I). These results demonstrate the successful independent recordings of two distinct neural populations in the ARC using Cre- or Flpo-dependent Ca^2+^ sensors. Our data also suggest a temporal succession characterized by a decrement in the ARC^Agrp^ neuron activity followed by the occurrence of SEs^kiss^ events.

### ARC^Agrp^ neurons negatively regulate SEs^kiss^ under a negative energy balance

If ARC^Agrp^ neurons are indeed involved in the modulation of SEs^kiss^, then manipulating their activities would have an impact on the frequency of SEs^kiss^. To test this hypothesis, we conducted a series of viral-genetic experiments. First, we chemogenetically activated ARC^Agrp^ neurons while monitoring SEs^kiss^ to investigate whether ARC^Agrp^ neurons could suppress SEs^kiss^ even after the removal of FR. Double heterozygous female *Kiss1-Cre; Agrp-Flpo* mice were subjected to the viral injection and FR. They were then administered an intraperitoneal injection of clozapine-*N*-oxide (CNO) and transferred to the AL condition (Figure 5A). Post-hoc histochemical analyses confirmed the targeting specificity of hM3Dq-mCherry (Figure 5B, C). In control mice that expressed only mCherry, SEs^kiss^ were frequently detected after removal of the FR, consistent with Figure 3G, even in the presence of CNO (Figure 5D–G). By contrast, chemogenetic activation of ARC^Agrp^ neurons in hM3Dq-expressing female mice significantly suppressed SEs^kiss^ within a 4-hour time window following CNO administration, during which CNO was effective (Figure 5D–G). Consistent with a preceding study^51^, the CNO-mediated activation of ARC^Agrp^ neurons resulted in increased body weight, indicating overeating and supporting the successful activation of ARC^Agrp^ neurons (Figure 5H). These data demonstrate that ARC^Agrp^ neurons negatively regulate the frequency of SEs^kiss^ during the recovery from FR.

**Figure 5.**
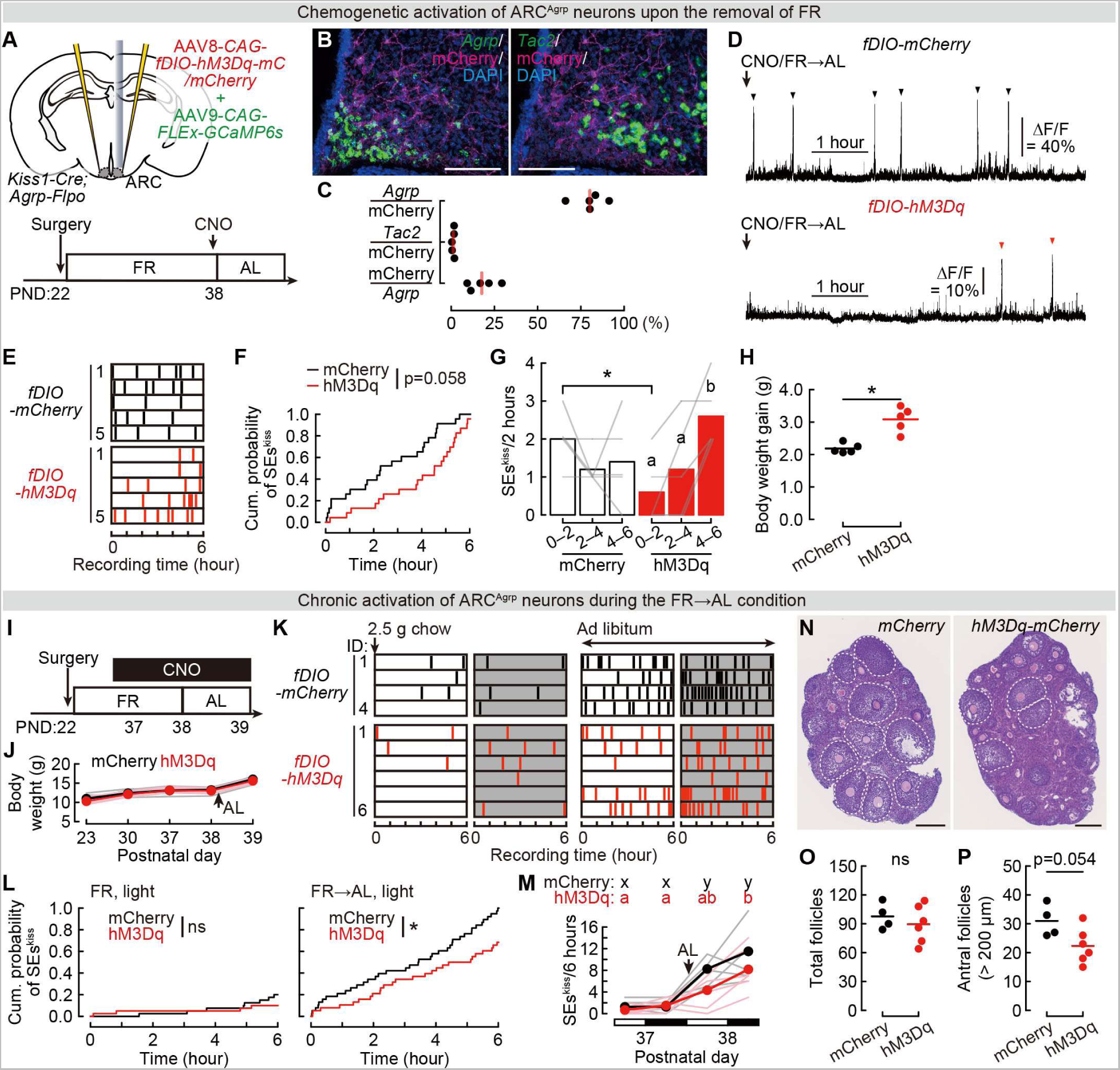
Peripubertal activation of ARC^Agrp^ neurons suppresses SEs^kiss^ upon removal of FR. (A) Schematic of the experimental design and time line. (B) Representative images of ARC showing *Agrp* mRNA expression (green) and hM3Dq-mCherry (magenta)(left), and *Tac2* mRNA expression (green, a marker of ARC^kiss^) and hM3Dq-mCherry (magenta)(right) counterstained with DAPI (blue). Scale bars, 100 μm. (C) Quantification of specificity (*Agrp*+/mCherry+), ratio of leaky expression (*Tac2*+/mCherry+), and efficiency (mCherry+/*Agrp+*) for the chemogenetic probe, n = 5. (D) Representative photometry traces showing SEs^kiss^ (arrowheads) following the removal of FR. Arrows indicate the timing of intraperitoneal administration of CNO and food supply. (E) Raster plots of SEs^kiss^ (vertical bars) in individual mCherry control (black) and hM3Dq-mCherry (hM3Dq, red) mice. (F) Cumulative probability of SE^kiss^ under the FR→AL conditions in the light period at PND 38. p = 0.058 by the Kolmogorov–Smirnov test. (G) Number of SEs^kiss^ per 2-hour time window following the CNO injection in the hM3Dq (red) and mCherry control (black) mice. Two-way ANOVA with repeated measures; AAV effect, ns; time course effect, p < 0.10; interaction effect, p < 0.01. *p < 0.05 by post-hoc two-sided unpaired *t*-test. Different letters (a, b) denote a significant difference (p < 0.05) by one-way ANOVA with repeated measures followed by the Tukey–Kramer post-hoc test. (H) Body weight gain of the hM3Dq (red) and mCherry control (black) mice between PND 38 and 39. *, p < 0.05 by the Mann–Whitney *U* test. n = 5 each for control and chemogenetic activation of ARC^Agrp^ neurons (E–H). (I) Experimental time line to investigate the phenotype on chronic activation of ARC^Agrp^. (J) Growth curve of the experimental animals. Two-way ANOVA with repeated measures; genotype effect, ns; PND effect, p < 0.01; interaction effect, ns. (K) Raster plots showing SEs^kiss^ (vertical bars) in mCherry control (black) and hM3Dq-mCherry (hM3Dq, red) mice. Open and gray boxes represent the light and dark periods, respectively. (L) Cumulative probability of SE^kiss^ under the FR and FR→AL conditions in the control (black) and the hM3Dq (red) mice. Total number of SEs^kiss^ in the control mice after removal of FR was normalized to 1. *, p < 0.05 by the Kolmogorov–Smirnov test. (M) Number of SEs^kiss^ per 6-hour time window. Two-way ANOVA with repeated measures; AAV effect, ns; period effect, p < 0.01; interaction effect, ns. Different letters (a, b, x, and y) denote a significant difference (p < 0.05) in the mCherry control (black) and hM3Dq (red) mice, respectively, by one-way ANOVA with repeated measures followed by the Tukey–Kramer post-hoc test. (N) Representative images of hematoxylin/eosin-stained ovarian sections from the mCherry control (black) and hM3Dq (red) mice one day after FR→AL (PND 39). Dotted circles indicate antral follicles (diameter, > 200 μm). Scale bars, 200 μm. (O, P) Total number of follicles (≥ secondary) (O) and antral follicles (P) in the mCherry control (black) and hM3Dq (red) mice. p-value by the Mann–Whitney *U* test. n = 4 for control and n = 6 for chronic activation of ARC^Agrp^ neurons (J–P). For more data, see Figure S5.

To examine the chronic impact of activating ARC^Agrp^ neurons on the ovarian phenotype, we administered CNO continuously via an osmotic pump from PND 37 to 39, with the FR removed at PND 38 (Figure 5I). While no differences were observed in body weight (Figure 5J), we observed a significant decrease in SEs^kiss^ frequency following the removal of FR (Figure 5K–M). CNO-treated hM3Dq-expressing female mice exhibited a trend of reduction in the number of antral follicles in the ovary, without altering the total number of follicles (Figure 5N–P). These data suggest that suppression of SEs^kiss^ by ARC^Agrp^ neurons during catch-up growth may negatively impact follicular development. It is noteworthy that under the chronic FR or AL condition, chemogenetic activation of ARC^Argp^ neurons did not affect ongoing SEs^kiss^ patterns (Figure 5L and Figure S5), implying that the activities of ARC^Agrp^ neurons impact only SEs^kiss^ during recovery from the negative energy balance.

To investigate the necessity of the functional ARC^Agrp^ neurons to suppress SEs^kiss^ under the negative energy balance, we conducted Ca^2+^ imaging of ARC^kiss^ neurons during PND 37–38 under the FR→AL condition, in combination with the use of a taCasp3-induced ablation approach^52^ to eliminate some ARC^Agrp^ neurons (Figure 6A). The mice consumed 2.5 grams of chow daily and maintained a body weight of approximately 13 grams under the FR condition (Figure 6B), despite a 40% reduction in the number of ARC^Agrp^ neurons (Figure 6C). The number of ARC^kiss^ neurons was not affected by this manipulation (Figure S6A, B), further supporting the specificity of our genetic targeting. In mice with ablated ARC^Agrp^ neurons, we noted a significantly elevated frequency of SEs^kiss^ during the light period upon food supply compared with control mice, whereas the frequency of SEs^kiss^ was comparable in the dark period when food availability was limited (Figure 6D–G). Upon removal of FR at PND 38, the mice with ablated ARC^Agrp^ neurons exhibited a significantly increased frequency of SEs^kiss^ (Figure 6F). A negative correlation trend was observed between the number of remaining ARC^Agrp^ neurons and the number of SEs^kiss^ (Figure S6C), further suggesting that ARC^Agrp^ neurons negatively impact SEs^kiss^.

**Figure 6.**
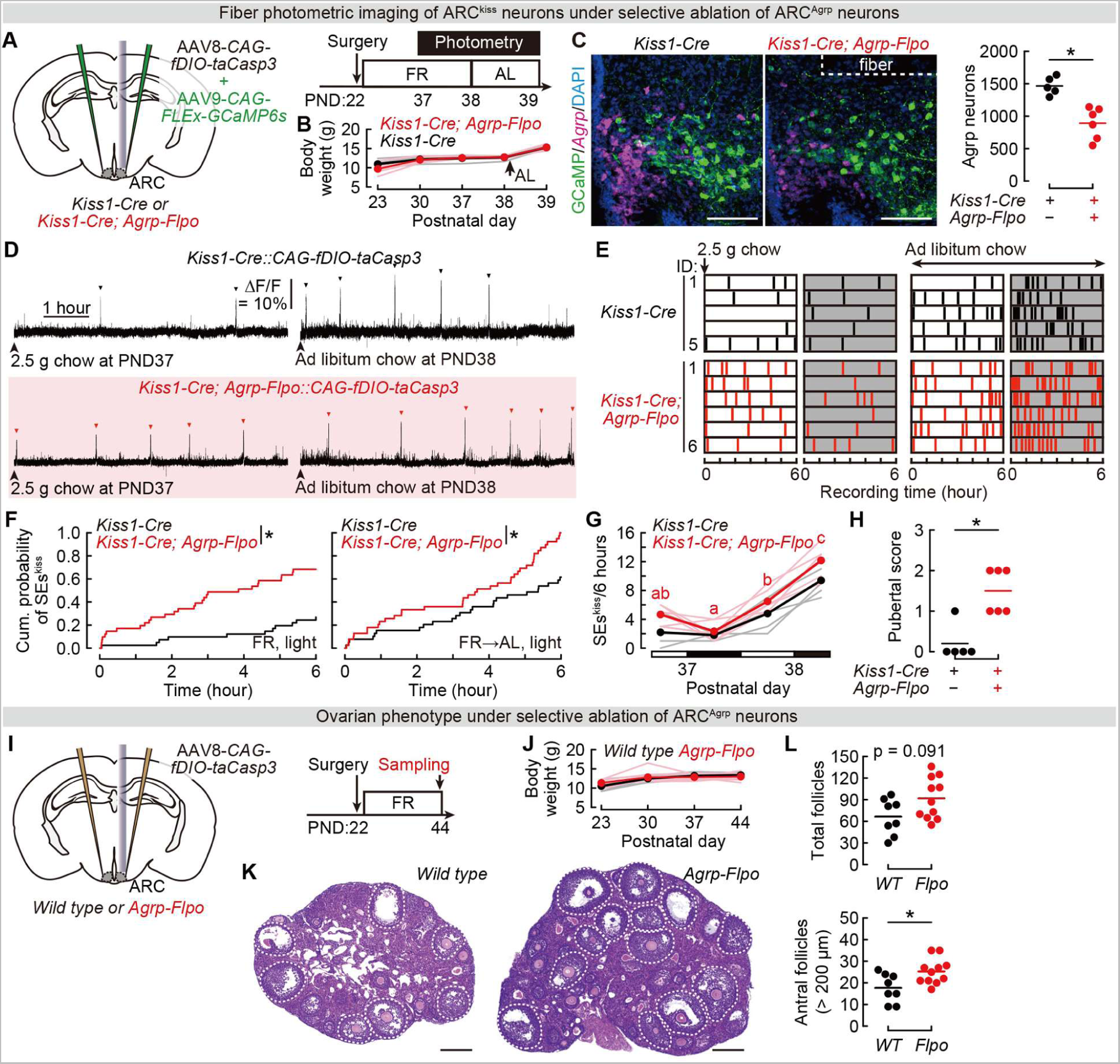
Peripubertal loss-of-function of ARC^Agrp^ neurons facilitates SEs^kiss^ under the FR condition. (A) Schematic of the experimental setup and time line. (B) Growth curve of the experimental animals. Two-way ANOVA with repeated measures; genotype effect, ns; PND effect, p < 0.01; interaction effect, p < 0.05. (C) Left: Representative sections of ARC showing *Agrp* mRNA expression (magenta) and GCaMP6s (green) counterstained with DAPI (blue). Scale bars, 100 μm. Right: Quantification of cell ablation efficiency. *, p < 0.05 by the Mann–Whitney *U* test. (D) Representative photometry traces showing SEs^kiss^ (arrowheads) during the light period under the FR (left) and after the removal of FR (right). Arrowheads indicate the timing of the food supply. (E) Raster plots showing SEs^kiss^ (vertical bars) in control (black) and ARC^Agrp^ neuron-ablated (red) mice. Open and gray boxes represent the light and dark periods, respectively. (F) Cumulative probability of SE^kiss^ under the FR and FR→AL conditions in the control (black) and ARC^Agrp^ neuron-ablation groups (red). Total number of SEs^kiss^ in *Kiss1*-*Cre*; *Agrp*-*Flpo* mice after removal of FR was normalized to 1. *, p < 0.05 by the Kolmogorov– Smirnov test. (G) Number of SEs^kiss^ per 6-hour time window. Two-way ANOVA with repeated measures; genotype effect, p < 0.05; period effect, p < 0.01; interaction effect, ns. Different letters (a–c) denote a significant difference (p < 0.05) in *Kiss1*-*Cre*; *Agrp*-*Flpo* mice by one-way ANOVA with repeated measures followed by the Tukey–Kramer post-hoc test. (H) Pubertal score one day after FR→AL (PND39), as in Fig. 1E. *, p < 0.05 by the Mann– Whitney *U* test. n = 5 for control and n = 6 for ablation of ARC^Agrp^ neurons (B–H). (I) Experimental time line to investigate the ovarian phenotype. (J) Growth curve of the *Agrp-Flpo* and wild-type control mice that received AAV *CAG-fDIO-taCasp3*. Two-way ANOVA with repeated measures; genotype effect, ns; PND effect, p < 0.01; interaction effect, p < 0.05. (K) Representative images of hematoxylin/eosin-stained ovarian sections from the *Agrp*-*Flpo* and wild-type mice at PND 44. Dotted circles indicate antral follicles (diameter, > 200 μm). Scale bars, 200 μm. (L) Total number of follicles (≥ secondary) (top) and antral follicles (bottom) in the *Agrp*-*Flpo* and wild-type mice. *, p < 0.05 by the Mann–Whitney *U* test. n = 8 for wild-type and n = 11 for *Agrp-Flpo* mice (J–L).

We also noticed that the pubertal score was significantly higher in ARC^Agrp^ neuron-ablated than in control mice (Figure 6H). This suggests that a loss of ARC^Agrp^ neurons promotes follicular development through an elevated frequency of SEs^kiss^. This view was supported by an additional cohort of ARC^Agrp^ neuron-ablation experiments in *Agrp-Flpo* mice under the chronic FR condition (Figure 6I). Although body weight was unchanged in ARC^Agrp^ neuron-ablated female mice (Figure 6J), H/E-stained sections showed more mature follicles (Figure 6K). Quantitative data showed that the total number of follicles tended to increase, with a significant increase of antral follicles (Figure 6L), suggesting irregular and partial sexual maturation. These data indicate that ARC^Agrp^ neurons are necessary for suppressing SEs^kiss^ and sexual maturation during the negative energy balance.

To probe more acute effects of suppressing ARC^Agrp^ neurons on SEs^kiss^, we employed hM4Di-mediated chemogenetic inhibition of ARC^Agrp^ neurons under chronic FR conditions (Figure S6D, E). At 2 hours after the CNO administration and the beginning of the imaging session, 2.5 grams of food were supplied. We observed a significant increase in SEs^kiss^ following food supply in CNO-administered hM4Di-expressing animals, whereas CNO had no apparent effect on SEs^kiss^ frequency before food supply (Figure S6F–I). Neither saline injection in hM4Di-expressing animals nor CNO administration in mCherry-expressing control animals affected the SEs^kiss^ pattern after food supply. A positive correlation was observed between the targeting efficiency of hM4Di to ARC^Agrp^ neurons and the number of SEs^kiss^ after CNO and food supply (Figure S6J). These findings indicate that the basal activities of ARC^Argp^ neurons after food supply are acutely required for suppressing SEs^kiss^ during the negative energy balance. Of note, an increased number of antral follicles was also supported in an additional cohort, wherein chemogenetic suppression of ARC^Agrp^ neurons was applied from PND 38 to 44 (Figure S6K–N). Collectively, our data demonstrate the indispensable role of ARC^Agrp^ neurons in suppressing SEs^kiss^ frequency and follicular maturation under chronic energy insufficiency.

## Discussion

Chronic negative energy balance, as observed in cases of malnutrition or anorexia nervosa, has been linked to a delayed onset of puberty^4^ and a subsequent decline in overall reproductive health later in life^53^. Despite growing interest in the molecular basis for the neuroendocrine control of puberty under normal and malnutrition conditions^1,54^, the temporal dynamics of SEs^kiss^ that are critical in regulating the hypothalamus–pituitary– gonad (HPG) axis^9,38,39^ remained elusive. Using fiber photometry, we elucidated the emergence, growth, and modulation of SEs^kiss^ during normal development and chronic energy deficiency. Importantly, the frequency of SEs^kiss^ was acutely modulated by dietary conditions through the specific hypothalamic neural circuit connecting feeding to reproductive centers. Here, we discuss the biological insights obtained by our study and its limitations.

Our data support a two-stage model of puberty initiation related to the temporal dynamics of SEs^kiss^ (Figure 7). The first stage (Stage I) is characterized by a low frequency of SEs^kiss^ (approximately 0.3 per hour), which occurs around PND 30 at a body weight of 12–14 g and triggers activation of the HPG axis, leading to vaginal opening. A 30% reduction in daily caloric intake persistently arrests animals in Stage I. Under normal feeding conditions, when the body weight surpasses 14 grams, animals proceed to the second stage (Stage II), which is characterized by a high frequency of SEs^kiss^ with 0.6–1.0 pulses per hour, supporting follicular growth to ovulatory follicles and ultimately leading to the first vaginal estrus. Upon the FR→AL transition, animals move into Step II within a few hours during catch-up growth. The activation of ARC^Agrp^ neurons appears to interfere with the transition from Stage I to Stage II (Figure 5). Conversely, inhibiting ARC^Agrp^ neurons allows animals under chronic FR conditions to progress partially to Stage II, increasing number of antral follicles (Figure 6). In this model, ARC^Agrp^ neurons function as one of the gatekeepers of the puberty checkpoint to prevent potentially harmful sexual maturation under the negative energy balance. Notably, our study does not preclude the potential involvement of alternative regulators, whether they be enhancers or inhibitors of SEs^kiss^ in response to the body’s metabolic status. In addition to our limited targeting efficiency of effector molecules to ARC^Agrp^ neurons, this could potentially account for why the chemogenetic stimulation of ARC^Agrp^ neurons in normally-fed animals did not result in alterations to SEs^kiss^ (Figure S5), and for why the inducing pubertal development was only partially successful via loss-of-function of ARC^Agrp^ neurons (Figures 6 and S6).

**Figure 7.**
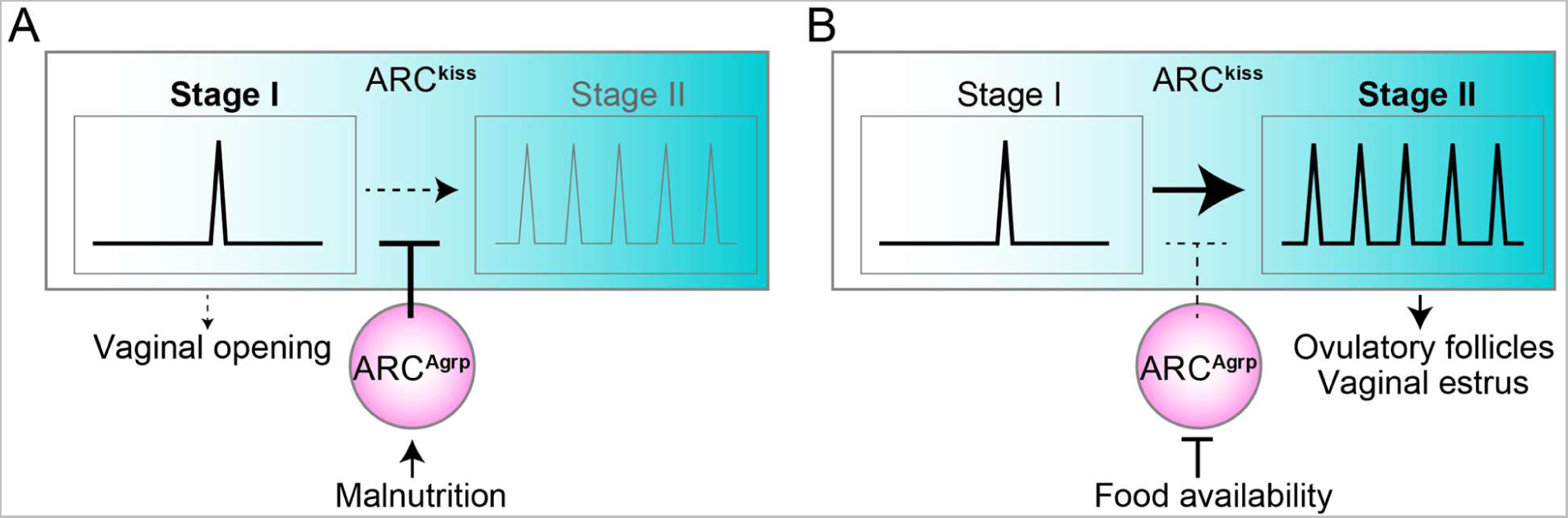
A two-stage model of the pubertal process. This model postulates that ARC^kiss^ neurons undergo two stages, characterized by low (~0.3 per hour) and high (0.6–1 per hour) frequencies of SEs^kiss^, and that the body’s energy balance in the peripubertal period can modulate the transition from Stage I to Stage II through the gatekeeping function of ARC^Agrp^ neurons. (A) Under malnourished conditions, such as those caused by FR, the activity of ARC^Agrp^ neurons is relatively high, leading to the suppression of ARC^kiss^ neurons. (B) When food availability improves (for example, by removing FR), disinhibition of ARC^Agrp^ neurons can facilitate SEs^kiss^ and promote the transition to Stage II. In this model, ARC^Agrp^ neurons serve as gatekeepers to the puberty checkpoint.

Our data established a strong correlation between decreased ARC^Agrp^ neuron activity, associated with feeding, and the occurrence of SEs^kiss^ (Figure 4). Furthermore, SEs^kiss^ frequency can be drastically altered within a few hours following changes in food availability (Figure 3). One potential benefit of this rapid response mechanism mediated by the ARC^Agrp^-to-ARC^kiss^ pathway is to adjust puberty timing in response to swiftly changing food resource conditions. Although leptin and blood glucose levels can effectively convey the body’s energy status to the brain^5,14^, hypoleptinemia and hypoglycemia are only observed in severely detrimental conditions. During food shortages, the decision to conserve energy for survival and food-seeking must be made well before humoral factors reach critical levels. In line with this perspective, our FR condition (a 30% reduction in daily food intake) did not induce hypoglycemia (Figure S2C) and minimally impacted plasma leptin levels (Figure 2F), despite a complete suppression of puberty onset as assessed by vaginal estrus (Figure 2C). The hunger signals represented by ARC^Agrp^ neuron activity^20^ allow for precise adjustments in reproductive investment based on daily dietary patterns. It is crucial to emphasize that our data do not disregard the importance of leptin signaling in the ARC^Agrp^ neurons^21^ or cellular energy sensors in the ARC^kiss^ neurons^4^ in the regulation of puberty. Instead, we posit that multiple regulatory systems, operating on different timescales, collectively ensure the robust control of puberty timing in response to changing dietary conditions.

The present study has several limitations. First, we did not uncover the neurotransmission mechanisms through which ARC^Agrp^ neurons hinder SEs^kiss^ during the negative energy balance. While ARC^Agrp^ neurons represent direct presynaptic GABAergic partners of ARC^kiss^ neurons^23^, they also express diverse neuropeptides that impact the activity and gene expression of downstream neurons^55,56^. To address this issue, future studies should combine pharmacology and cell-type-specific gene knockouts. It is also important to discriminate the mono- and multi-synaptic pathways originating from the ARC^Agrp^ neurons to affect SEs^kiss^. Second, we did not examine the molecular underpinnings behind the development and modulation of SEs^kiss^. SEs^kiss^ are shaped by the coordinated cell-autonomous action of neurokinin B and dynorphin A^6^, with the frequency modulated by diverse factors ranging from estrogen signaling to inflammatory stress^57^. Future studies, such as single-cell RNAseq of ARC^kiss^ neurons, could potentially uncover how the molecular foundation of SEs^kiss^ may be modulated under various conditions, including those in the negative energy balance. Finally, whether the ARC^Agrp^-to-ARC^kiss^ pathway contributes to the food-linked regulation of reproductive functions in both mature females and males represents an important open question. Despite the deceleration of the estrus cycle in normally-nourished adult female mice following chemogenetic stimulation of ARC^Agrp^ neurons^23^, the potential involvement of ARC^kiss^ neurons in effecting this outcome still remains ambiguous (see Figure S5). Our established suite of imaging and manipulation toolkits will facilitate molecular and circuit-based investigations into the complex interaction between feeding and reproductive capabilities under various conditions, including youth, adulthood, good health, and pathological states.

## Acknowledgments

We thank the staff at the RIKEN Biosystems Dynamics Research animal facility for animal care and *in vitro* fertilization, Hiroko Tsukamura (Nagoya Univ), Tomomi Karigo (Johns Hopkins University), Ken Murata (the University of Tokyo), and members of the Miyamichi laboratory for the critical reading of the manuscript, Manami Fujii and Gen-ichi Tasaka for the technical assistance, and the University of North Carolina Vector Core and Canadian Neurophotonics Platform Viral Vector Core Facility for the AAV productions. This study was supported by Grants-in-Aid from the Japan Society for the Promotion of Science (JSPS) to T.G. (Nos. 19K16270 and 21K15194) and K.M. (Nos. 20K20589 and 21H02587), Mitsubishi foundation and a Takeda Life Science grant to K.M., and a RIKEN internal fund to K.M. T.G. was also supported by RIKEN Special Postdoctoral Researchers Program.

## Author contributions

T.G. and K.M. conceived the experiments. T.G. performed the experiments and analyzed the data with technical support by M.H. and S.I. *Agrp-Flpo* mice were generated by M.H., T.A., and H.K. T.G. and K.M. wrote the paper with contributions from all co-authors.

## Declaration of Interests

The authors declare that they have no competing interests.

## Supplementary Figures

**Figure S1.**
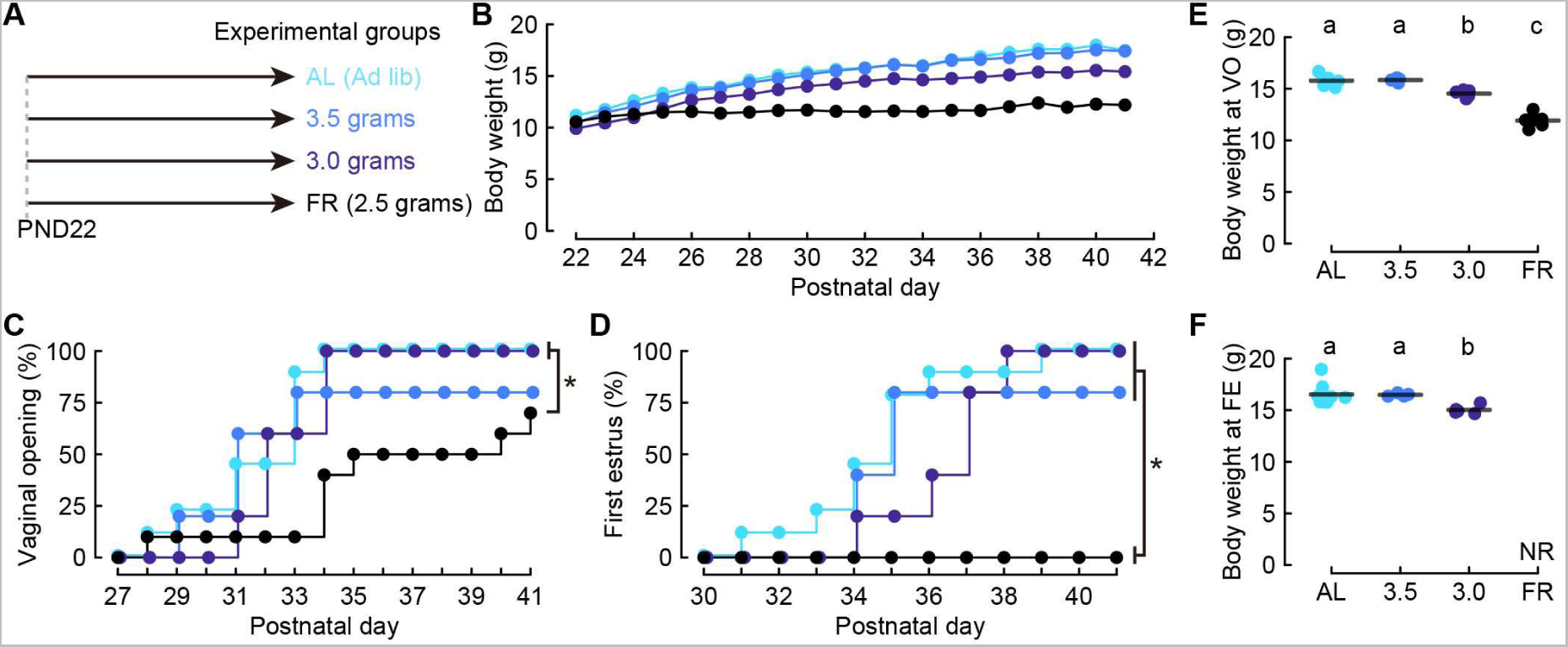
Puberty onset under different daily food amounts, related to Figure 1. (A) Timetable of food conditions. PND, postnatal day. (B) Average growth curves of the peripubertal mice under the following four conditions: i) AL (light blue, the same as AL in Figure 1B); ii) 3.5 grams/day food supply (blue); iii) 3.0 grams/day food supply (navy); iv) 2.5 grams/day food supply (black), equivalent to the FR condition in Figure 1B. (C, D) Cumulative probability of puberty onset, measured by vaginal opening (C) and vaginal cytological first estrus (D), in the peripubertal mice under the above four conditions. *, p < 0.05 by the Kolmogorov–Smirnov test with Bonferroni correction. The data for the AL and FR groups correspond to Figure 1C. (E, F) Body weight at vaginal opening (E) and first estrus (F). Letters (a–c) denote a significant difference (p < 0.05) by one-way ANOVA followed by the Tukey–Kramer post-hoc test. NR, no record. n = 9 for AL, n = 5 for 3.5 grams/day, n = 5 for 3.0 grams/day, and n = 10 for FR (B–F).

**Figure S2.**
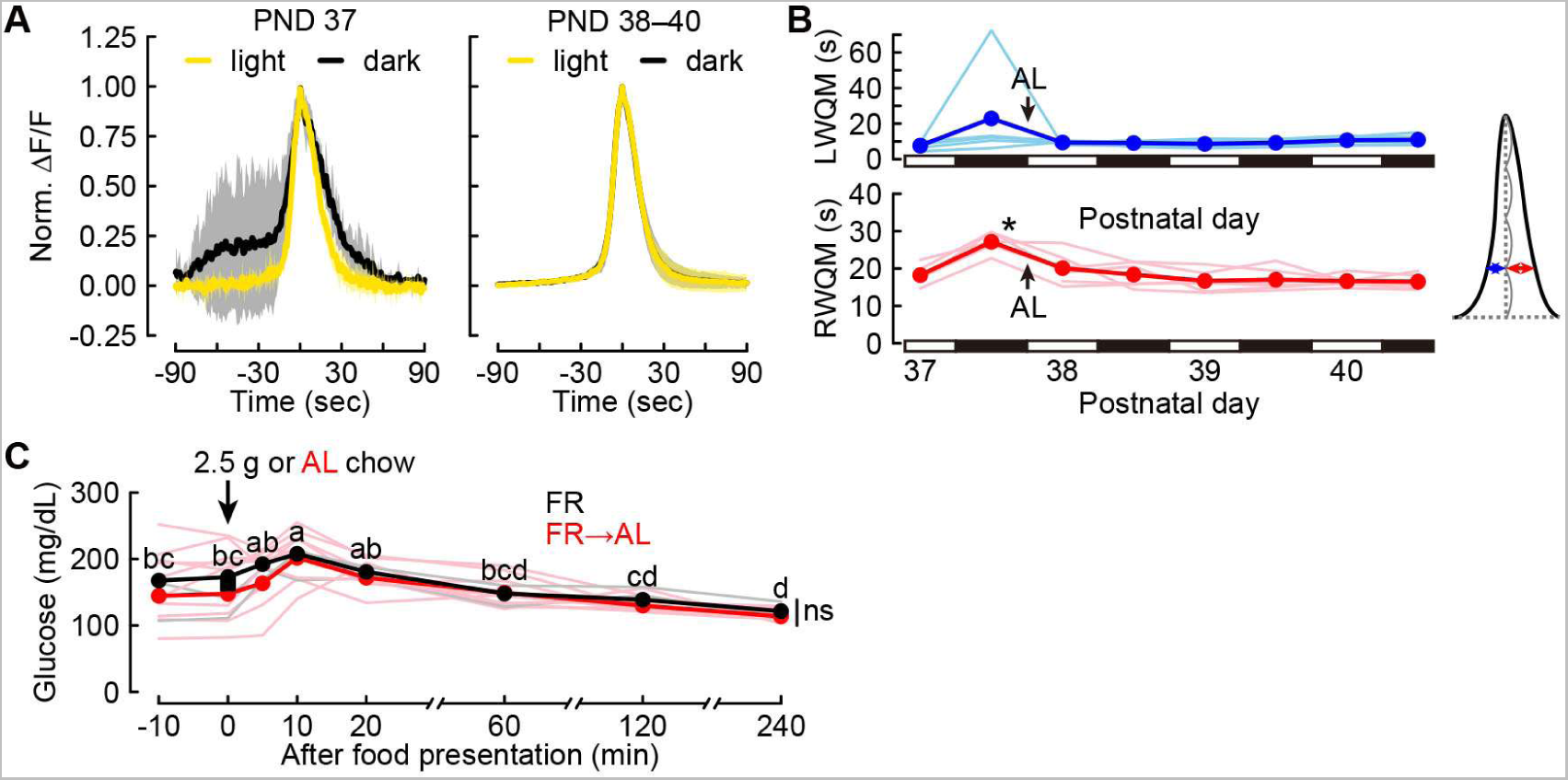
Additional data regarding peripubertal SEs^kiss^, related to Figure 3. (A) Waveform of SE^kiss^ under the FR condition (PND 37, left) and after FR removal (PND 38–40, right) in the FR→AL group. Yellow and black lines show the averaged photometry traces in the light (soon after food supply) and dark (> 12 hours after food supply) periods, respectively, with shadows representing standard deviations. (B) LWQM and RWQM of SE^kiss^ at PND 37–40 in the FR→AL group. *, p < 0.05 by one-way ANOVA with repeated measures followed by the Tukey–Kramer post-hoc test, n = 6. (C) Blood glucose levels under the FR condition (black circles) and soon after FR removal (red circles) in the FR→AL group, n = 6. No FR control mice (n = 4) are indicated by black squares at time point 0. Two-way ANOVA with repeated measures; feeding condition effect, ns; time course effect, p < 0.01; interaction effect, ns. Different letters (a–d) denote a significant difference (p < 0.05) by one-way ANOVA with repeated measures followed by the Tukey–Kramer post-hoc test.

**Figure S3.**
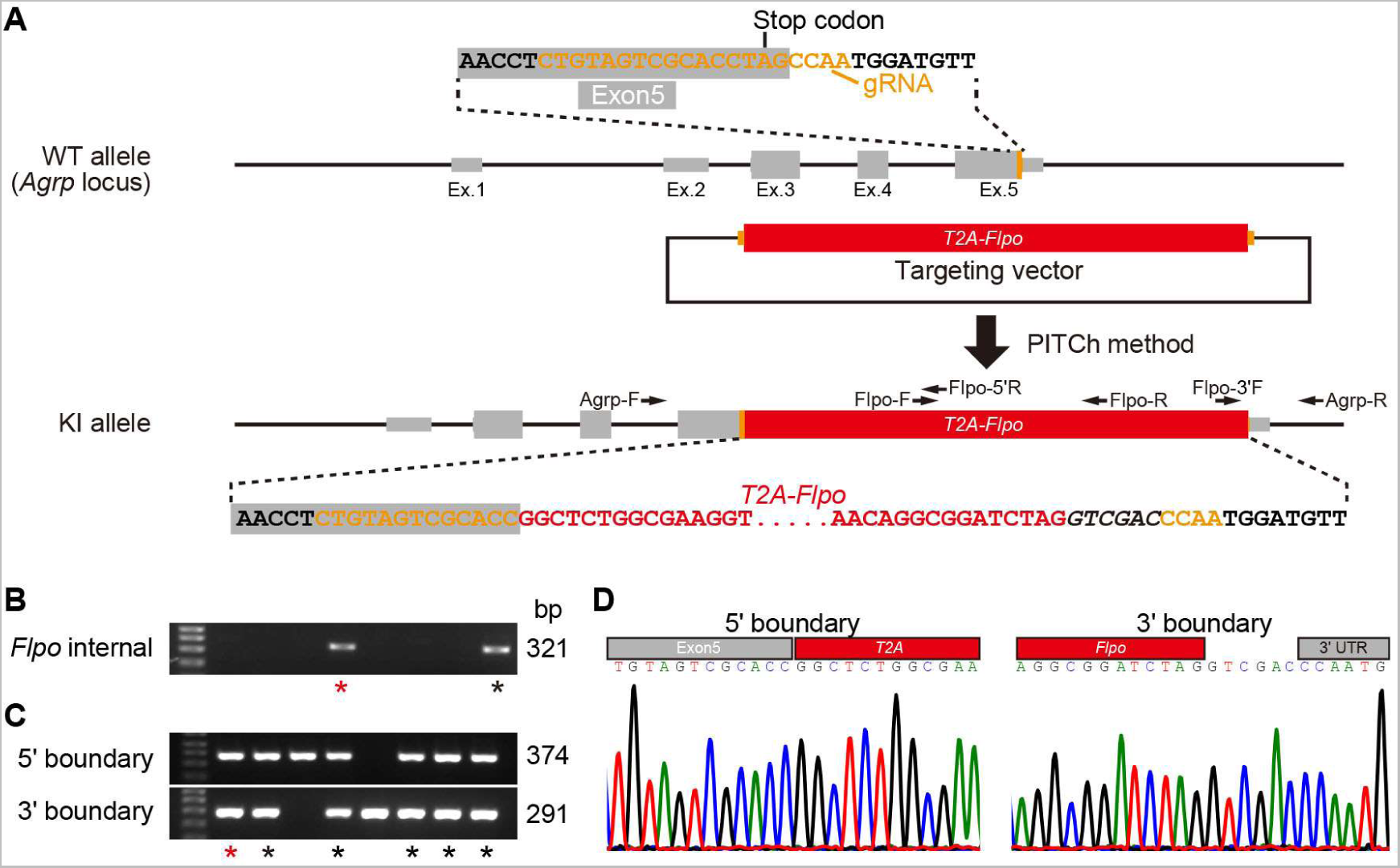
Generation of *Agrp-Flpo* mice, related to Figure 4. (A) Illustration of the knock-in strategy using the PITCh method ^58^. A *T2A-Flpo* cassette was inserted into the coding end of the *Agrp* locus. (B, C) Screening of *Agrp-Flpo* founder mice. Genotyping was performed using an internal *Flpo* sequence (B) and the 5’ and 3’ boundaries (C). Black asterisks indicate candidates for F_0_ founders by PCR screening. Red asterisks indicate the established F_0_ lines following sequence analysis. The following PCR primers were used: Flpo-F: Flpo-R for Flpo internal sequence, Agrp-F: Flpo-5’R for the 5’ boundary, and Flpo-3’F: Agrp-R for the 3’ boundary. The size of the PCR product is indicated at the right of the gel images. See the Methods section for sequence information. (D) Confirmation of the 5’ and 3’ boundary sequences.

**Figure S4.**
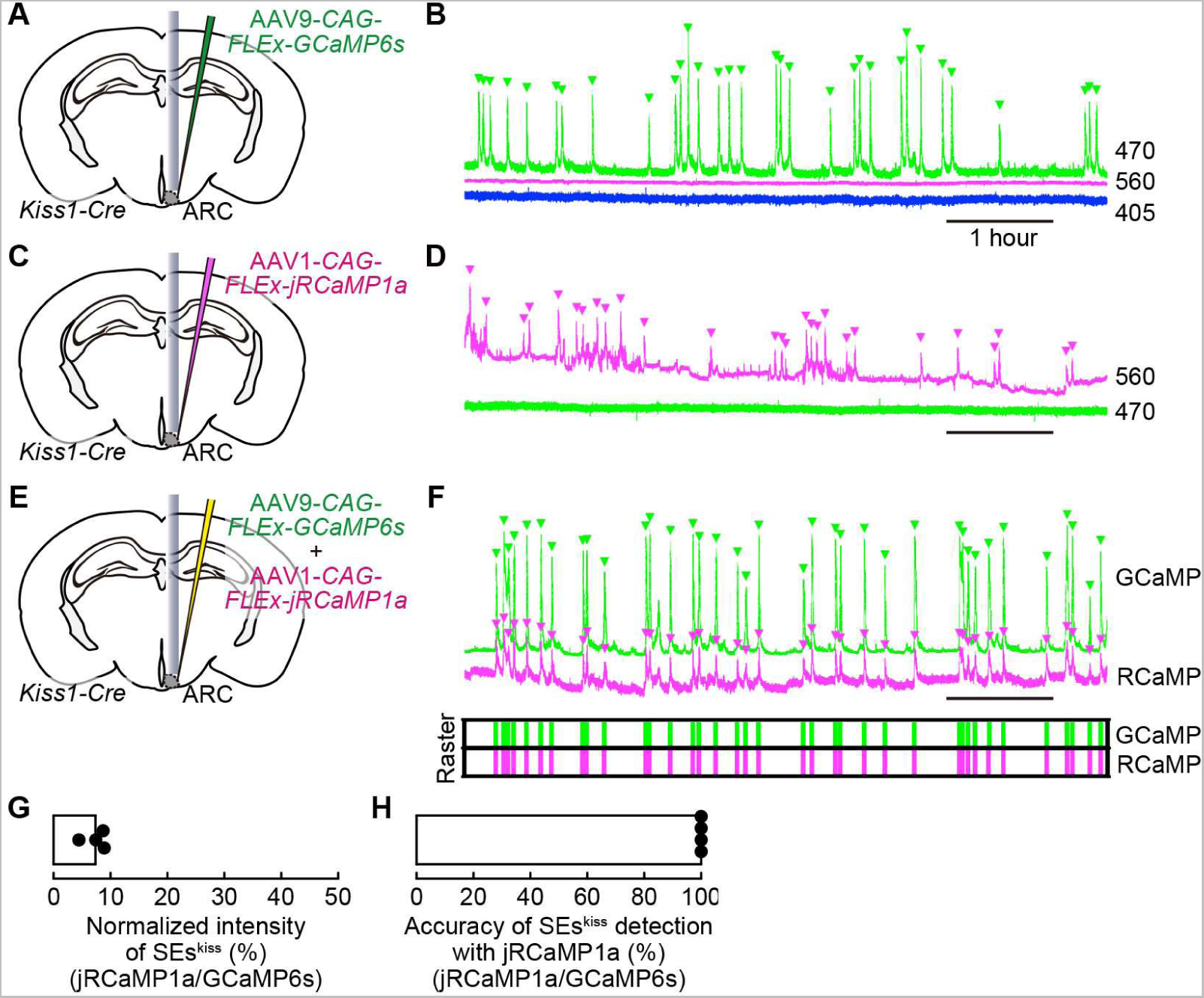
Control experiments for dual-color fiber photometry, related to Figure 4. (A, C, E) Schematic of the experimental settings. In *Kiss1-Cre* female mice, the AAV carrying a Cre-dependent GCaMP6s (A), the AAV carrying a Cre-dependent jRCaMP1a (C), or both AAVs (E) were injected into the ARC. Ovariectomy was performed to eliminate the negative feedback signals from the gonads for better visualization of SEs^kiss^. An optical fiber was inserted above the ARC for fiber photometry. (B, D, F) Representative photometry traces of the 470-nm channel (green, Ca^2+^-dependent GCaMP6s signals), 560-nm channel (magenta, Ca^2+^-dependent jRCaMP1a signals), and 405-nm channel (blue, non-Ca^2+^ dependent signals of GCaMP6s, shown only in panel B). Green and magenta arrowheads indicate individual SEs^kiss^. The raster plots at the bottom of panel F show SEs^kiss^ as vertical bars, which demonstrate the precise 1:1 correspondence of SEs^kiss^ detected by GCaMP6s and jRCaMP1a. (G) Quantification of the normalized intensities of SEs^kiss^ (jRCaMP1a/GCaMP6s) detected in animals that received both AAVs. The intensities of jRCaMP1a signals are < 10% of those detected with GCaMP6s. (H) Ratio of SEs^kiss^ detected with GCaMP6s that were also detected with jRCaMP1a in animals that received both AAVs. n = 4 (panels G and H).

**Figure S5.**
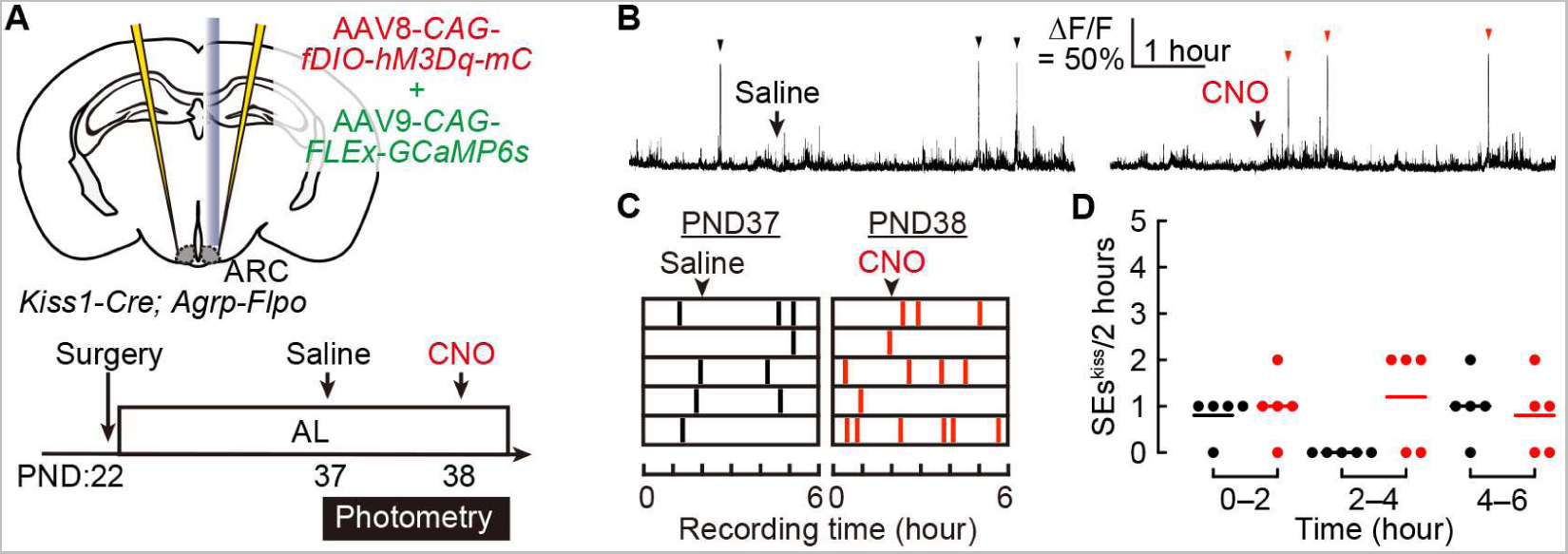
Peripubertal chemogenetic activation of ARC^Agrp^ neurons under the AL condition does not affect SEs^kiss^, related to Figure 5. (A) Schematic of the experimental paradigm and time line to assess the effect of chemogenetic activation of ARC^Agrp^ neurons on SEs^kiss^ under the AL condition. (B) Representative photometry traces showing SEs^kiss^ (black and red arrowheads) in the ovary-intact double heterozygous *Kiss1-Cre*; *Agrp-Flpo* female mice. Arrows indicate the time of intraperitoneal injection of saline or CNO. (C, D) Raster plots of SEs^kiss^ (vertical bars) (C) and number of SEs^kiss^ per 2-hour time window (D) in the hM3Dq-expressing mice treated with saline (black) or CNO (red). No significant difference was found by two-way ANOVA with repeated measures.

**Figure S6.**
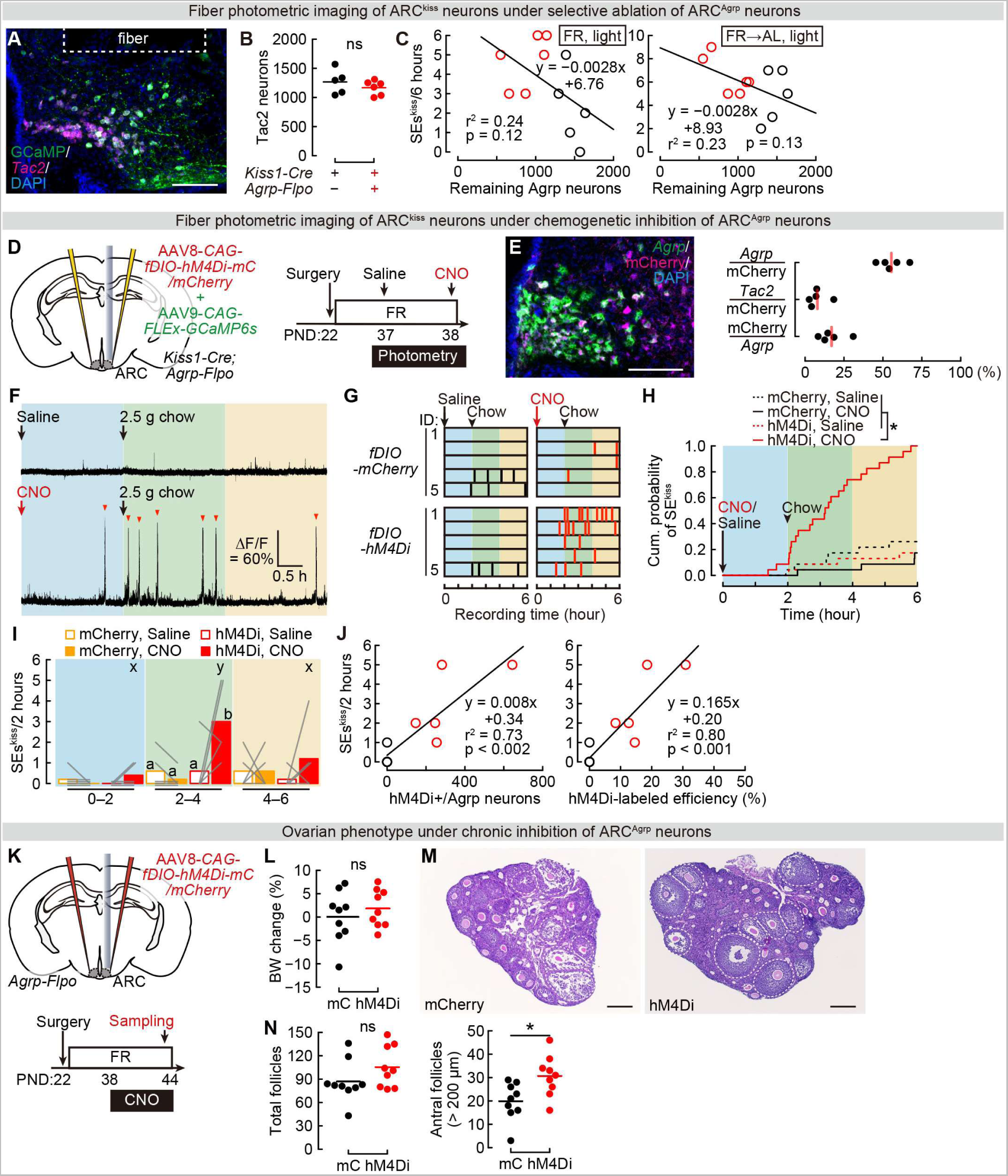
Additional data regarding the loss-of-function experiments of ARC^Agrp^ neurons, related to Figure 6. (A) Representative ARC section showing *Tac2* mRNA expression (magenta, a marker of ARC^kiss^ neurons) and GCaMP6s (green) counterstained with DAPI (blue) after imaging from ARC^Agrp^ neuron-ablated (*Kiss1-Cre*; *Agrp-Flpo*) mice. Scale bar, 100 μm. (B) Quantification of nonspecific ablation (number of *Tac2*+ cells). No statistical difference was found by the Mann–Whitney *U* test. (C) Weak negative correlations between the remaining *Agrp* neurons (the X-axis) and the numbers of SEs^kiss^ in the light period under the FR condition (PND 37) or the FR→AL condition (PDN38) (the Y-axis). Black and red circles represent data from the control and ARC^Agrp^ neuron-ablation groups, respectively. Black lines, a line fit. n = 5 for control and n = 6 for ablation of ARC^Agrp^ neurons (B, C). (D) Schematic of the experimental setup and time line for chemogenetic inhibition of ARC^Agrp^ neurons. CNO was administered immediately before the initiation of photometry imaging at PND 38, with a saline control session conducted 1 day earlier. (E) Left: Representative ARC section showing *Agrp* mRNA expression (green) and hM4Di-mCherry (magenta) counterstained with DAPI (blue). Scale bar, 100 μm. Right: Quantification of specificity (*Agrp*+/mCherry+), ratio of leaky expression (*Tac2*+/mCherry+), and efficiency (mCherry+/*Agrp*+), n = 5. (F) Representative photometry traces showing SEs^kiss^ (arrowheads) during the light period. Arrows indicate the timing of the intraperitoneal administration of saline or CNO and food supply. (G) Raster plots of SEs^kiss^ (vertical bars) in individual mice injected with saline (black) or CNO (red). (H) Cumulative probability of SE^kiss^ in mCherry control (black) and hM4Di-expressing (red) mice injected with saline (dotted lines) or CNO (solid lines). *, p < 0.05 by the Kolmogorov–Smirnov test with Bonferroni correction. (I) Number of SEs^kiss^ per 2-hour time window in mCherry control (orange) and hM4Di-expressing (red) mice injected with saline (open boxes) or CNO (closed boxes). Because hM4Di-expressing mice with CNO differed significantly from the other groups, we performed a statistical test of time within the group and comparisons between groups at 2–4-hour windows. Two-way ANOVA with repeated measures; AAV effect, p < 0.05; drug effect, ns, interaction effect, p < 0.05. Different letters (a and b) denote a significant difference (p < 0.05) by one-way ANOVA followed by the Tukey–Kramer post-hoc test. Different letters (x and y) denote a significant difference (p < 0.05) by one-way ANOVA with repeated measures followed by the Tukey–Kramer post-hoc test. (J) Correlation between the targeting efficiency of hM4Di to ARC^Agrp^ neurons (left, number of mCherry+ *Agrp*+ neurons per ARC; right, ratio of mCherry+/Agrp+) and the numbers of SEs^kiss^ per 2-hour time window following food supply. Red and black circles represent data from the hM4Di-expressing animals and mCherry-expressing control animals, respectively. The p-values were calculated based on the null hypothesis assuming no correlation. Black lines, a line fit. n = 5 each for mCherry control and hM4Di-expressing mic (G–J). (K) The experimental time line to investigate the ovarian phenotype. CNO was subcutaneously administrated with an osmotic pump during PND 38–44. (L) Change in body weight (BW) (%) at PND 44 relative to PND 38 in the mCherry (mC, black) control and hM4Di-expressing (hM4Di, red) mice. (M) Representative images of hematoxylin/eosin-stained ovarian sections from the mCherry and hM4Di mice at PND 44. Dotted circles indicate antral follicles (diameter, > 200 μm). Scale bars, 200 μm. (N) Total number of follicles (≥ secondary) (left) and antral follicles (right) in the mCherry control and hM4Di-expressing female mice. ns, not significant, and *, p < 0.05 by the Mann–Whitney *U* test in (L and N). n = 9 each for mCherry control and hM4Di-expressing mice (L–N).

**Movie S1. Representative traces of dual-color fiber photometry illustrating food supply effects, related to Figure 4**.

Dual-color Ca^2+^ imaging conducted in a PND 38 *Kiss1-Cre*; *Agrp-Flpo* double heterozygous female. Top, a side view of the cage. Bottom, fiber photometry traces of jRCaMP1a (magenta, ARC^kiss^ neurons) and GCaMP6s (green, ARC^Agrp^ neurons). Approximately 10 seconds in this movie, food is supplied, leading to a decline in GCaMP6s signals. Around the 29-second mark, the subject initiates feeding, soon accompanied by a SE^kiss^ detected by jRCaMP1a signals. The movie is shown at 4 × real-time speed.

## RESOURCE AVAILABILITY

### Lead contact

Further information and requests for resources and reagents should be directed to and will be fulfilled by the Lead Contact, Kazunari Miyamichi.

### Materials availability

All data are available in the main paper and supplementary materials. All materials, including the *Agrp-Flpo* mice and plasmids, are available through requests to the corresponding authors.

### Data and code availability

- Data reported in this paper are available from the lead contact upon reasonable request.
- All custom analysis codes are available from the lead contract upon reasonable request.
- Any additional information required to reanalyze the data reported in this paper is available from the Lead Contact upon request.

## EXPERIMENTAL MODEL AND SUBJECT DETAILS

All animal experiments were approved by the Institutional Animal Care and Use Committee of the RIKEN Kobe Branch. C57BL/6J female mice were purchased from Japan SLC, Inc. (Shizuoka, Japan). *Kiss1-Cre* (Jax #017701) and *Ai162* (*TIT2L-GC6s-ICL-tTA2*, Jax #031562)*-D* mice were purchased from the Jackson Laboratory. *Agrp-Flpo* mice were originally generated in this study (described below in detail). Animals were maintained at the animal facility of the RIKEN Center for Biosystems Dynamics Research under an ambient temperature (18–23 °C) and 12-hour light, 12-hour dark cycle schedule. The mice were allowed ad libitum access to a laboratory diet (MFG; Oriental Yeast, Shiga, Japan; 3.57 kcal/g) and water unless otherwise mentioned.

## METHOD DETAILS

### Generation of *Agrp-Flpo* knock-in mice

The *Agrp-Flpo* knock-in mouse line (Accession No. CDB0162E: https://large.riken.jp/distribution/mutant-list.html) was generated by CRISPR/Cas9-mediated knock-in in zygotes, as previously described^49^. The SV40 nuclear localization signal (NLS) was added to the 5’ of *Flpo* open reading frame by PCR primers 5’-GGCGCGCCACCATGGCTCCTAAGAAGAAGAGGAAGGTGATGAGCCAGTTCGACAT CCTG; 5’-GTCGACTCAGATCCGCCTGTTGATGTAG. The plasmid *pTCAV-FLEx(loxP)-FlpO* (Addgene #67829) was used as a PCR template. Then, the targeting vector consisting of *T2A-Flpo* was constructed by the PCR-based method and inserted just before the stop codon of *Agrp*. The guide RNA (gRNA) sites were designed by using CRISPRdirect^59^ to target the upstream and downstream locations of the stop codon (Figure S3). For microinjection, a mixture of two CRISPR RNAs (crRNAs) (50 ng/μL), trans-activating crRNA (tracrRNA) (200 ng/μL), donor vector (10 ng/μL), and Cas9 protein (100 ng/μL) was injected into the pronucleus of a C57BL/6 one-cell stage zygote. Agrp crRNA (5’-CUG UAG UCG CAC CUA GCC AAg uuu uag agc uau gcu guu uug), PITCh 3 crRNA (5’-GCA UCG UAC GCG UAC GUG UUg uuu uag agc uau gcu guu uug), and tracrRNA (5’-AAA CAG CAU AGC AAG UUA AAA UAA GGC UAG UCC GUU AUC AAC UUG AAA AAG UGG CAC CGA GUC GGU GCU) were purchased from FASMAC (Atsugi, Japan). As a result, 70 F_0_ founder mice were obtained, 14 (6 males and 8 females) of which were targeted as identified by 5’ and 3’ junction PCR. We further analyzed 5’ and 3’ junction and full-length sequencing from three males. The following primers were used for the PCR and sequence analysis.

For the detection of the *Flpo* internal sequence: Flpo-F 5’-CTGGCCACATTCATCAACTGCGG; Flpo-R 5’-CTTCTTCAGGGCCTTGTTGTAGCTG.

For the 5’ boundary: Agrp-F 5’-TGACCTCAGTCCACTGCCACCCTAC; Flpo-5’R 5’-TCTTGCACAGGATGTCGAACTGGCTC.

For the 3’ boundary: FLPo-3’F 5’-AGCATCAGATACCCCGCCTGGAACG; Agrp-R 5’-GCCTGGTGCCTTAAACTCGCCCATATA.

For the full-length: Agrp-F 5’-TGACCTCAGTCCACTGCCACCCTAC; Agrp-R 5’-GCCTGGTGCCTTAAACTCGCCCATATA.

The PCR products using the primer sets were subcloned into the pCR Blunt II TOPO vector for each (Zero Blunt TOPO PCR Cloning Kit, Thermo Fisher) and sequenced using M13-Foward and M13-Reverse primers. The germline transmission of the *Agrp-Flpo* allele was confirmed by genotyping of F_1_ mice. The line was established by two males that harbored a targeted sequence identified by PCR and sequencing. Genotyping PCR was performed using the *Flpo* internal primers Flpo-F and Flpo-R, as mentioned above (see Figure S3A for details).

### Estrus cycle monitoring by vaginal cytology

The peripubertal mice were carefully checked for the state of vaginal opening. If the mice exhibited vaginal opening, 10 μL of tap water was gently pipetted once or twice into the vagina with minimum insertion to prevent the induction of pseudopregnancy. These vaginal smears were placed on a slide glass and evaluated for the types of cells as described below. The stage of the estrus cycle was determined based on the presence or absence of leukocytes and cornified and nucleated epithelial cells, in accordance with previous studies^41,42^. In Figures 2E and 6H, the peripubertal mice were assessed on a 0–3 scale according to their vaginal state: absence of vaginal opening (score 0); the presence of vaginal opening and diestrus (1), proestrus (2), and estrus (3).

### Viral preparations

The following AAV vectors were purchased from Addgene. The titer is shown as genome particles (gp) per milliliter.

AAV serotype 9 *CAG-FLEx-GCaMP6s* (1.5 × 10^13^ gp/mL, #100842-AAV9)

AAV serotype 1 *CAG-FLEx-NES-jRCaMP1a-WPRE-SV40* (1.4 × 10^13^ gp/mL, #100846-AAV1)

AAV serotype 8 *Ef1a-fDIO-mCherry-WPREpA* (1.8 × 10^13^ gp/mL, #154867-AAV8)

The following AAV vectors were purchased from the Canadian Neurophotonics Platform Viral Vector Core Facility (RRID: SCR_016477).

AAV serotype 9 *CAG-fDIO-GCaMP6s* (6.9 × 10^12^ gp/mL, construct-1347-aav2-9)

AAV serotype 8 *hSyn-fDIO-hM3D(Gq)-mCherry* (4.7 × 10^12^ gp/mL, construct-1457-aav2-8)

AAV serotype 8 *hSyn-fDIO-hM4D(Gi)-mCherry* (6.8 × 10^12^ gp/mL, construct-1243-aav2-8)

To generate *hSyn-fDIO-taCasp3*, the taCasp3-TEVp sequence from pAAV *FLEx-taCasp3-TEVp* (Addgene, #45580) was subcloned into the pAAV *hSyn-fDIO-WPREpA* sequence from pAAV *hSyn-fDIO-hM3D(Gq)-mCherry-WPREpA* (Addgene, #154868). The AAV vector (AAV serotype 8 *hSyn-fDIO-taCasp3*, 5.7 × 10^11^ gp/mL) was generated by the UNC viral core using the plasmid pAAV *hSyn-fDIO-taCasp3*, which has been deposited to Addgene (#207997).

### Stereotaxic injection

For the injection of AAV and an optical fiber at postnatal days 21–22, the mice were anesthetized with an intraperitoneal injection of saline containing 65 mg/kg ketamine (Daiichi Sankyo, Tokyo, Japan) and 13 mg/kg xylazine (X1251; Sigma-Aldrich), and head-fixed to a stereotaxic apparatus (#68045, RWD).

To perform the experiments in Figure 1, for prepubertal *Kiss1-Cre; Ai162* female mice, an optical fiber (NA = 0.50, core diameter = 400 μm from Kyocera, Tokyo, Japan) was placed over the ARC using the following coordinates from the bregma: posterior 1.8 mm, lateral 0.2 mm, and ventral 5.7 mm defined on a brain atlas^60^. The injected fiber was fixed on the skull with dental cement (SanMedical, Shiga, Japan). The postsurgical mice were housed individually and allowed ad libitum access to a laboratory diet (MFG; Oriental Yeast) and water.

To perform the experiments in Figure 3, the AAV carrying a Cre-dependent GCaMP6s (AAV9 *CAG-FLEx-GCaMP6s-WPRE-SV40*) was injected into the ARC of prepubertal *Kiss1-Cre* female mice using the following coordinates from the bregma: posterior 1.8 mm, lateral 0.2 mm, and ventral 5.8 mm. In Figure 4, a 1:1 mixture of the AAV carrying a Cre-dependent jRCaMP1a (AAV1 *CAG-FLEx-NES-jRCaMP1a-WPRE-SV40*) and an AAV carrying a Flpo-dependent GCaMP6s (AAV9 *CAG-fDIO-GCaMP6s*) was injected into the ARC of peripubertal *Kiss1-Cre; Agrp-Flpo* female mice. In Figures 5 and S5, a 1:3 mixture of the AAV carrying a Cre-dependent GCaMP6s and an AAV carrying a Flpo-dependent hM3D(Gq)-mCherry (AAV8 *hSyn-fDIO-hM3D(Gq)-mCherry*), or a 1:1 mixture of the AAV carrying a Cre-dependent GCaMP6s and an AAV carrying a Flpo-dependent mCherry (AAV8 *Ef1a-fDIO-mCherry-WPREpA*), was injected into the ARC of *Kiss1-Cre; Agrp-Flpo* mice. In the first part of Figure 6, a 1:4 mixture of an AAV carrying a Cre-dependent GCaMP6s and an AAV carrying a Flpo-dependent taCasp (AAV8 *hSyn-fDIO-taCasp3-WPREpA*) was injected into the ARC of prepubertal *Kiss1-Cre; Agrp-Flpo* or *Kiss1-Cre* female mice. In the second part of Figure 6, the AAV carrying a Flpo-dependent taCasp was injected into the ARC of prepubertal *Agrp-Flpo* or wild-type female mice. In the second part of Figure S6, a 1:3 mixture of the AAV carrying a Cre-dependent GCaMP6s and an AAV carrying a Flpo-dependent hM4D(Gi)-mCherry (AAV8 *hSyn-fDIO-hM4D(Gi)-mCherry*), or a 1:1 mixture of an AAV carrying a Cre-dependent GCaMP6s and an AAV carrying a Flpo-dependent mCherry, was injected into the ARC of *Kiss1-Cre; Agrp-Flpo* female mice. In the last part of Figure S6, the AAV carrying a Flpo-dependent hM4D(Gi)-mCherry, or an AAV carrying a Flpo-dependent mCherry, was injected into the ARC of *Agrp-Flpo* female mice. A total of 200 nL of AAV was injected into the ARC at a speed of 50 nL/minute using a glass capillary regulated by a micro syringe pump injector (UMP3; World Precision Instruments, Sarasota, FL, USA) in all experiments. Soon after AAV injection, an optical fiber (NA = 0.50, core diameter = 400 μm from Kyocera) was placed above the ARC using the following coordinates from the bregma: posterior 1.8 mm, lateral 0.2 mm, and ventral 5.7 mm defined on a brain atlas^60^. The injected fiber was fixed on the skull with dental cement. The postsurgical mice were housed individually and allowed ad libitum access to food (baby food; CLEA Japan, Tokyo, Japan; 3.92 kcal/g) and water until the body weight reached 10.5 grams.

To perform the experiments in Figure S4, the AAV carrying a Cre-dependent GCaMP6s, an AAV carrying a Cre-dependent jRCaMP1a, or a 1:1 mixture of an AAV carrying a Cre-dependent GCaMP6s and an AAV carrying a Cre-dependent jRCaMP1, was injected into the ARC of ovariectomized *Kiss1-Cre* female mice using the following coordinates from the bregma: posterior 2.0 mm, lateral 0.2 mm, and ventral 5.9 mm. An optical fiber (NA = 0.50, core diameter = 400 µm from Kyocera) was placed above the ARC using the following coordinates from the bregma: posterior 2.0 mm, lateral 0.2 mm, and ventral 5.8 mm.

### Fiber photometry recording

Fluorescence signals were acquired using a fiber photometry system based on a previously published design^30^. All the optical components were purchased from Doric Lenses (Quebec, Canada). We performed chronic Ca^2+^ imaging by delivering excitation light (470 nm modulated at 530.481 Hz) and collecting emitted fluorescence using an integrated Fluorescence Mini Cube (Doric Lenses, iFMC4_IE(405)_E(460–490)_F(500– 550)_S). Light collection, filtering, and demodulation were performed using the Doric photometry setup and Doric Neuroscience Studio Software (Doric Lenses). The 470-nm signal was recorded as calcium-dependent GCaMP6s. The power output at the tip of the fiber was about 4–10 μW. The signals were initially acquired at 12 kHz and then decimated to 120 Hz. In addition, we performed dual-color Ca^2+^ imaging by delivering excitation light (470 nm modulated at 530.481 Hz, 560 nm modulated at 369.549 Hz) and collecting emitted fluorescence using the integrated Fluorescence Mini Cube (Doric Lenses, iFMC6_IE(400-410)_E1(460–490)_F1(500–540)_E2(555–570)_F2(580–680)_S). The 470-nm and 560-nm signals were recorded as calcium-dependent GCaMP6s and jRCaMP1a, respectively. The power output at the tip of the fiber was about 24–25 μW. Ca^2+^ imaging was performed twice per day for 6 hours each in the light and dark periods, with 6-hour inter-imaging intervals. The imaging was performed with mice in which SE^kiss^ had been detected on the day before or 2 days before the imaging.

For the analyses, we used a homemade R code. Because the signal intensity of SEs^kiss^ varies widely due to variations in surgery, we first calculated the average ΔF/F height of the stereotyped SE^kiss^ peak of each mouse from several visually obvious peaks with the full width at half-maximum threshold over 10 seconds^30^. The ΔF/F was calculated by (F_t_ – F_0_) / F_0_, where F_t_ is the recorded signal at time = *t* and F_0_ is the average of signals over the entire 6 hours of recording. Then, SEs^kiss^ were automatically detected by using the findpeaks function in the R project, with the peak height threshold as 0.4-fold of the average ΔF/F height. To analyze the reliable peak height and waveform further (Figures 3H and S2), we extracted the raw data from –240 s to +120 seconds when the peak was set to 0 seconds. We then recalculated the ΔF/F by (F_t_ – F_0_) / F_0_, where F_t_ is the recorded signal at time = *t* and F_0_ is the average of signals from –240 s to –120 s. Left half-width at quarter maximum (LWQM) and right half-width at quarter maximum (RWQM) were determined by the recalculated ΔF/F value (Figure S2B). To show the averaged waveform (Figure S2A), the recalculated ΔF/F data were downsized at 10 Hz.

In Figure 1, Ca^2+^ imaging was monitored every 2–4 days for PND 24–45 along with body weight and vaginal opening. An activity heatmap of ARC^Agrp^ neurons (Figure 4E) is represented as Z-scores calculated based on a 15- to 5-minute time window prior to food supply for each animal. Cross-correlation between the peak of SEs^kiss^ and Z score of the activity of ARC^Agrp^ neurons (Figure 4G) was calculated using ccf function in R project. In Figures 3 and 5, Ca^2+^ signals were monitored along with body weight. In Figure 6A–H, Ca^2+^ imaging was monitored along with body weight and vaginal opening.

### Food restriction

When the mice weighed more than 10.5 grams by PND 23, the food supply was removed and the bedding was renewed. The mice were then supplied with 2.50 ± 0.01 grams of baby food (CLEA Japan Inc., Tokyo, Japan; 3.92 kcal/g) every day at Zeitgeber time 4–6 under ad libitum access to water. In Figure S1, mice were also supplied with 3.00 ± 0.01 or 3.50 ± 0.01 grams of baby food for comparison. The mice were manually or automatically supplied the amount of food at 24-hour intervals through a customized feeding system using an automatic feeder for pets (highshop, #FC0196200pufF) fixed to the cage top.

### Chemogenetic manipulation

Clozapine *N*-oxide (CNO, #4936, Tocris bioscience) was dissolved at 0.5 mg/mL for intraperitoneal injection and 6.1 mg/mL for osmotic pump, respectively. CNO was intraperitoneally injected to achieve a dose of 2 mg/kg. Under isoflurane anesthesia, an Alzet osmotic minipump (model 1007D; flow rate 0.5 μL/hour, 100 μL capacity, Durect, Cupertino, CA, USA) filled with 6.1 mg/mL CNO was surgically implanted into the subcutaneous cavity of the mice in the evening on postnatal day 36. The animals (average weight, 12–14 grams) received CNO at approximately 5 mg/kg/day under this condition.

### Blood glucose monitoring

Blood glucose levels were monitored using a blood glucose meter for laboratory animals (SUGL-001; ForaCare Japan, Tokyo, Japan) after obtaining blood samples from the tail. The animals were habituated for 2 days before the measurements to prevent potential stress-induced blood glucose elevation.

### Plasma leptin measurement

Leptin supplementation was administered via an Alzet osmotic minipump (model 1002; flow rate 0.25 μL/hour, 100 μL capacity, Durect). The pump was filled with 7.5 mg/mL leptin (498-OB; R&D Systems, Inc., Minneapolis, MN, USA) and surgically implanted into the subcutaneous cavity of the food-restricted mice in the evening on postnatal day 23.

Plasma leptin levels were measured using an enzyme-linked immunosorbent assay (Morinaga Institute of Biological Science, Yokohama, Japan) using blood samples obtained from the tail in the morning according to the manufacturer’s instructions.

### Histology

Mice were anesthetized with isoflurane and perfused with phosphate-buffered saline (PBS) followed by 4% paraformaldehyde (PFA) in PBS. The brain and ovaries were post-fixed with 4% PFA in PBS overnight. Thirty-micrometer coronal brain sections (every fourth section) were made using a cryostat (Leica). To generate cRNA probes for *Tac2* and *Agrp*, DNA templates were amplified by PCR from the whole-brain cDNA (Genostaff, cat#MD-01). A T3 RNA polymerase recognition site (5’-*AATTAACCCTCACTAAAGGG*) was added to the 3’ end of the reverse primers. The forward (F) and reverse (R) primers to generate DNA templates for cRNA probes are as follows: F 5’-*AGCCAGCTCCCTGATCCT*; R 5’-*TTGCTATGGGGTTGAGGC* for *Tac2*, and F 5’-*CCCAAGAATGGACTGAGCAT*; R 5’-*TGCGACTACAGAGGTTCGTG* for *Agrp*. DNA templates for *Tac2* and *Agrp* were subjected to *in vitro* transcription with DIG (cat#11277073910)- and fluorescein (cat#11685619910)-RNA labeling mix and T3 RNA polymerase (cat#11031163001) according to the manufacturer’s instructions (Roche Applied Science).

Fluorescent *in situ* hybridization (ISH) combined with anti-GFP immunohistochemical staining was performed as previously reported^61^. In brief, after hybridization and washing, brain sections were incubated with peroxidase (POD)-conjugated anti-fluorescein (Roche Applied Science, cat#11426346910, 1:500) antibody overnight at 4 °C. On the next day, signals were amplified with TSA-plus Cyanine 5 (NEL745001KT, Akoya Biosciences; 1:70 in 1× plus amplification diluent) for 25 minutes. After washing with PBS containing 0.1% Tween-20 (PBST) for 5 minutes, POD was inactivated with 2% sodium azide in PBS for 15 minutes, followed by 10 minutes of washing three times with PBST. The sections were then incubated with horseradish peroxidase (HRP)-conjugated anti-DIG (Roche Applied Science cat#11207733910, 1:500) and anti-GFP (Aves Labs cat#GFP-1010, 1:200) antibodies overnight. Signals were amplified by TSA-plus Cyanine 3 (NEL744001KT, Akoya Biosciences; 1:70 in 1× plus amplification diluent) for 25 minutes, followed by washing, and then GFP-positive cells were visualized by anti-chicken Alexa Fluor 488 (Jackson Immuno Research cat#703-545-155, 1:250). PBS containing 50 ng/mL 4’,6-diamidino-2-phenylindole dihydrochloride (DAPI; Sigma-Aldrich, cat#D8417) was used for counter nuclear staining. Images were acquired using an Olympus BX53 microscope equipped with a 10× (N.A. 0.4) objective lens. Cells were counted manually.

Fluorescent ISH combined with anti-mCherry immunohistochemical staining was performed as follows. After hybridization and washing, brain sections were incubated with POD-conjugated anti-fluorescein (1:500) antibody overnight at 4 °C. On the next day, signals were amplified with TSA-plus Cyanine 5 for 25 minutes. After washing with PBST for 5 minutes, POD was inactivated with 2% sodium azide in PBS for 15 minutes, followed by 10 minutes of washing three times with PBST. The sections were then incubated with HRP-conjugated anti-DIG (1:500) antibody overnight and treated with TSA-plus biotin (NEL749A001KT; Akoya Biosciences), followed by streptavidin-Alexa Fluor 488 (S32354, Invitrogen). After 10 minutes of washing three times with PBST, the sections were incubated with goat anti-mCherry (Acris Antibodies cat#AB0040-200, 1:400) antibody overnight, followed by washing, and then mCherry-positive cells were visualized by anti-goat Alexa 555 (Sigma cat#A-32816, 1:250). PBS containing 50 ng/mL DAPI was used for counter-nuclear staining. Images were acquired using an Olympus BX53 microscope equipped with a 10× (N.A. 0.4) objective lens. Cells were counted manually.

To detect jRCaMP1a signals, before fluorescent ISH, brain sections were mounted on cover glass using Fluoromount (Diagnostic BioSystems cat#K024) and images were acquired using an Olympus BX53 microscope. After the cover glass was slipped out in the PBS bath, fluorescent ISH combined with anti-GFP immunohistochemical staining was performed as described above.

We compared *Tac2* gene expression in the ARC as a robust marker of ARC^kiss^ neurons because the expression level of the *Kiss1* gene is not sufficient for stable detection by conventional ISH.

The ovaries were embedded in paraffin. Serial 5-µm sections (every 10^th^) were made using a microtome. Paraffin sections were deparaffinized with xylene, hydrated with ethanol, and HE-stained for light microscopy. Follicle size was quantified by Fiji software.

### Statistical analysis

The statistical details of each experiment, including the statistical tests used, the exact value of n, and what n represents, are described in each figure legend. We used R version 4.3.0 (http://www.R-project.org/) for all analyses. Differences at p < 0.05 were considered significant.

## Notes

### Competing Interest Statement

The authors have declared no competing interest.

